# Clonal landscape of human nephrons

**DOI:** 10.64898/2026.11.19.763808

**Authors:** Kosuke Ieiri, Nobuyuki Kakiuchi, Koichi Watanabe, Tomonori Hirano, Shun Kawaguchi, Hirona Maeda, Yoshikage Inoue, Tomomi Nishimura, Taisho Noda, Masanori Motomura, Tomoe Nakagawa, Masahiro M. Nakagawa, Ryunosuke Saiki, Yotaro Ochi, Satoru Miyano, Yoshinao Oda, Ryota Nakagawa, Tatsuki R. Kataoka, Marin Takaso, Masahito Hitosugi, Genshiro Fukuchi, Dai Takamatsu, Shohei Nagakawa, Tokiyoshi Tanegashima, Shigehiro Tsukahara, Jun Mutaguchi, Satoshi Kobayashi, Shunsuke Goto, Takashi Matsumoto, Masaki Shiota, Masatoshi Eto, Seishi Ogawa

**Affiliations:** Department of Pathology and Tumor Biology, Graduate School of Medicine, Kyoto University, Kyoto, Japan; Department of Urology, Graduate School of Medical Sciences, Kyushu University, Fukuoka, Japan; The Hakubi Center for Advanced Research, Kyoto University, Kyoto, Japan; Department of Gastroenterology and Hepatology, Graduate School of Medicine, Kyoto University, Kyoto, Japan; Department of Gynecology and Obstetrics, Graduate School of Medicine, Kyoto University, Kyoto, Japan; Department of Diagnostic Pathology, Kyoto University Hospital, Kyoto, Japan; Department of Gastrointestinal Surgery, Kyoto University Hospital, Kyoto, Japan; Department of Surgery, Japanese Red Cross Fukui Hospital, Fukui, Japan; Department of Urology, Graduate School of Medicine, The University of Tokyo, Tokyo, Japan; Department of Integrated Analytics, M&D Data Science Center, Institute of Integrated Research, Institute of Science Tokyo, Tokyo, Japan; Department of Anatomic Pathology, Pathological Sciences, Graduate School of Medical Sciences, Kyushu University, Fukuoka, Japan; Department of Pathology, Iwate Medical University, Iwate, Japan; Department of Diagnostic Pathology, Shiga General Hospital, Shiga, Japan; Department of Legal Medicine, Shiga University of Medical Science, Shiga, Japan; Department of Urology, Saiseikai Yahata General Hospital, Fukuoka, Japan; Department of Urology, Saga-ken Medical Centre Koseikan, Saga, Japan; Department of Urology, Fukuoka Central Hospital, Fukuoka, Japan; School of Medicine, International University of Health and Welfare, Fukuoka, Japan; Institute for the Advanced Study of Human Biology (WPI-ASHBi), Kyoto University, Kyoto, Japan; Department of Innovative Medicine, Faculty of Medicine, Kindai University, Osaka, Japan

## Abstract

The nephron is a highly compartmentalized structure, where distinct patterns of somatic mosaicism may be established and contribute to altered kidney functions with aging. Here we show extensive remodeling of aged nephrons by mosaic clones using integrated sequencing of >1,000 microdissected samples collected from anatomically distinct nephron segments. Striking differences were observed across segments. Proximal tubules exhibited markedly higher mutation burdens with mutational signatures characteristic of clear cell renal cell carcinoma and larger clonal expansion than other segments. Cancer-related mutations were observed only infrequently and largely limited to proximal tubules, while copy number abnormalities were quite common, most frequently involving chromosomes 18, 10, 7, X, 22, 3 and 9, in a segment-specific manner. Clone size appears to depend more on cell turnover than on driver events. Particularly, in collecting ducts, numerous clones independently acquiring chromosome 18 loss evolved over years and ultimately replaced 15-47% of collecting duct cells in aged donors despite minimal increase in their clone size, likely reflecting strong fitness advantage of chromosome 18-deficient clones under slow tissue turnover. Chromosome 18-deficient cells significantly correlated with fibrosis and showed reduced expression of chromosome 18 genes, including those involved in sodium handling (*NEDD4L*) and TGF-β signaling (*SMAD2/4/7*). In line with this, GSEA of chromosome 18-deficient cells revealed enhanced hallmark pathways overlapping those associated with acute kidney injury that are related to TGF-β signaling and epithelial-mesenchymal transition, and inflammatory responses. These findings illustrate how nephrons are remodeled with aging, providing insights into the role of somatic remodelling in altered kidney structures and functions with aging.

---

During the past decade, studies on early cancer evolution in non-neoplastic tissues have revealed frequent remodeling of aged or inflamed tissues by somatic mosaicism typically comprising many clones carrying cancer-related driver mutations^1–7^. Widely observed in most human tissues, such remodeling by somatic clones appears to be an almost inevitable consequence of aging and also affected by tissue-specific factors, such as mutational processes, cell turnover rates, selective pressures imposed by microenvironment, and tissue architecture^1,4,5,8–11^. Despite their strong association with common cancer-related mutations, mosaic clones may not necessarily be destined for cancer development but may contribute to non-neoplastic human diseases or compromised organ functions in the elderly, which has been intensively studied in clonal hematopoiesis^12,13^, but poorly investigated in the context of somatic mosaicism in solid tissues.

The kidney consists of numerous functional units, nephrons, with approximately one million nephrons per organ^14^. The nephron is a highly compartmentalized, closed structure and is divided into distinct functional segments, including the glomerulus, proximal and distal tubules, and the loop of Henle in between, which ultimately connects to the collecting duct network^15^ (Fig. 1a). In the context of somatic mosaicism, this closed structure may impose strict spatial constraints on clonal expansion and therefore influence clonal dynamics, while the unique ecosystems surrounding nephrons may dictate distinct patterns of clonal evolution and the establishment of somatic mosaicism. Nephrons operate under low oxygen tension and are exposed to a steep osmotic gradient maintained along the corticomedullary axis and also to high concentrations of diverse chemicals and metabolites with mutagenic potential, which renal tubular cells reabsorb from glomerular filtrates or actively transport from the circulation, particularly in proximal segments^16–18^. In addition, with aging, kidneys undergo progressive structural and functional changes characterized by chronic inflammation, enhanced TGF-β signaling, interferon responses, and proinflammatory cytokine signaling, and progressive fibrosis, which have been implicated in age-related renal dysfunction and chronic kidney disease (CKD)^19,20^. Conversely, these age-related alterations may shape the selective pressures acting on somatic clones within nephron epithelium.

**Figure 1.**
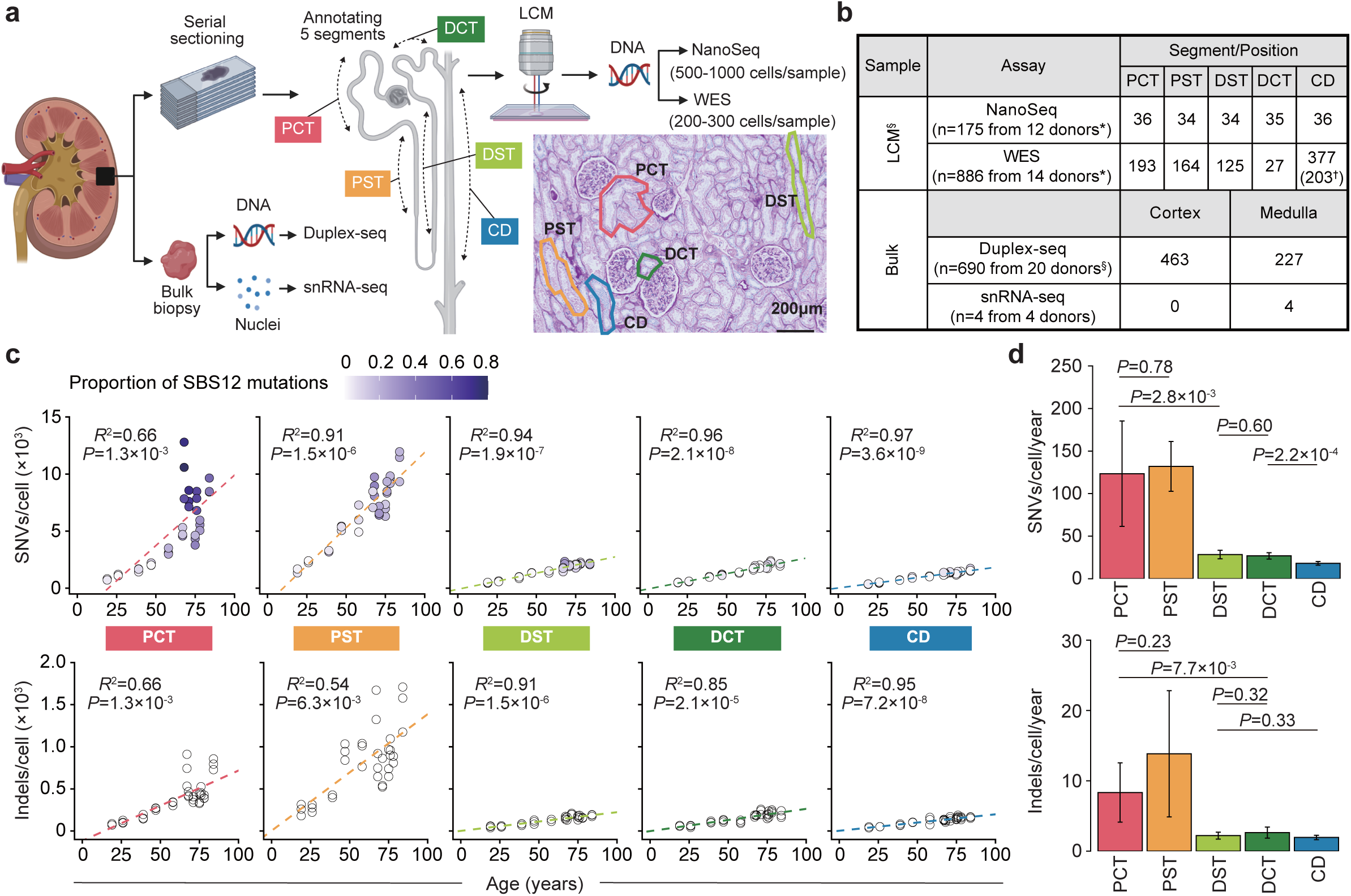
Study design and mutation accumulation in normal nephrons. a, Laser-capture microdissection (LCM) was performed to collect micro-scale samples from five distinct nephron segments including PCT, PST, DST, DCT and CD, followed by NanoSeq or WES. In parallel, bulk renal tissues were also analyzed using Duplex-seq and snRNA-seq. **b,** Numbers of donors and samples in each assay. *Seven and §11 donors are overlapped between assays. ^†^Samples independently collected for validation experiments (STable 1). **c,** Correlation between mean numbers of SNVs and indels/cell and donor age in indicated nephron segments. Content of SBS12 mutations is shown as a color gradient. *R*^2^ and *P* values were obtained using a linear regression model. **d,** Mean numbers (±95% confidence interval) of SNVs and indels/cell/year. *P* values were calculated based on the interaction term of the linear regression model.

In the context of carcinogenesis, on the other hand, nephron epithelium is the origin of different types of renal cancers, including clear cell renal cell carcinoma (ccRCC), papillary renal cell carcinoma (pRCC), and chromophobe renal cell carcinoma (chRCC)^21,22^. Thus, it is of particular interest to investigate somatic mosaicism in aged nephrons across the distinct functional segments to gain mechanistic insight into the cellular origin of RCCs and also into the declining renal function with aging^23^. Unfortunately, however, due to the complex architecture of nephrons and the technical challenges in isolating each nephron segment separately, somatic mosaicism within the distinct kidney nephrons has not been explored.

In this study, we performed extensive sequencing analysis of micro-scale samples isolated from distinct nephron segments in different age groups using laser-capture microdissection (LCM). Somatic mosaicism in each segment was characterised by assessing segment-specific mutational burdens and their signatures, RCC-related driver mutations, copy number abnormalities (CNAs), and expansion of mosaic clones. Through these analyses, we revealed drastic remodeling of kidney nephrons with aging, particularly in collecting ducts, driven by segment-specific somatic mosaicism. The functional consequences of the remodeling were further investigated using single-nucleus RNA sequencing of collecting duct cells.

## Somatic mutations in normal nephrons

In total, we collected 1,061 LCM samples from five anatomically distinct nephron segments from 19 Japanese donors with or without renal tumors (Fig. 1b, STable 1): proximal convoluted and straight tubules (PCT and PST), distal straight and convoluted tubules (DST and DCT), as well as collecting duct (CD). These distinct segments were recognized by their characteristic staining patterns and tubular morphology based on hematoxylin and eosin (H&E) and periodic acid–Schiff (PAS) staining (Fig. 1a; Methods), where the high accuracy of assignment (>95.8% for each segment) was validated by immunohistochemistry using segment-specific antibodies (ED_Fig. 1). Among these, 175 samples from 12 donors aged 19 to 84 years were analyzed using ultra-low-error-rate sequencing (NanoSeq^24^) to evaluate mutational burdens in distinct nephron segments, while the remaining 886 (consisting of 683 + extended 203 samples) were used to evaluate positively selected mutations and CNAs as well as clonal expansion, using whole exome sequencing (WES) (Fig. 1a-b, STable 1-2).

In NanoSeq, randomly reading 0.97x whole genome on average (range: 0.14x-2.54x) at a very low error rate (∼5×10^-9^), we were able to obtain an accurate estimate of mutation accumulation rates even from small polyclonal cell populations (STable 3). Each LCM sample for NanoSeq was generated by combining 3-4 microdissected areas from the same nephron segment type, comprising a total of ∼1,000 cells. Across all segments, both single-nucleotide variants (SNVs) and insertions and deletions (indels) linearly increased with age (Fig. 1c). However, the mutation accumulation rate varied markedly across segments (Fig. 1d, STable 4): PCT and PST showed notably higher accumulation rates than those thus far reported in normal tissues^1,25^ (123 and 132 SNVs and 8.3 and 13.8 indels/cell/year, respectively), compared with DST (28.4 SNVs and 2.2 indels/cell/year) and DCT (26.8 SNVs and 2.6 indels/cell/year). CDs showed the lowest accumulation rates (18.1 SNVs and 2.0 indels/cell/year). Compared with other segments, proximal tubules also exhibited greater inter-individual variability in mutational burden, suggesting differences in proximal tubule-specific mutational processes across donors. To understand the mutational processes underlying these variations, we next analyzed mutational signatures across different nephron segments.

### Mutational signatures in normal nephrons

In total, 196,238 SNVs and 19,894 indels detected in 175 LCM samples were subjected to mutational signature analysis. Hierarchical Dirichlet Process (HDP)-based deconvolution analysis using COSMIC v3.4 signatures as priors identified seven SNV signatures showing very high cosine similarity (CS) to COSMIC SBS1, SBS5, SBS8, SBS12, and SBS40a–c, and therefore they were designated as these COSMIC labels hereafter (Fig. 2a; Methods). In agreement with a previous report^26,27^, an additional analysis with SBS-192 trinucleotide context (Methods) revealed moderate to strong transcriptional strand bias for SBS40b and SBS40c, and SBS12 (ED_Fig. 2a).

**Figure 2.**
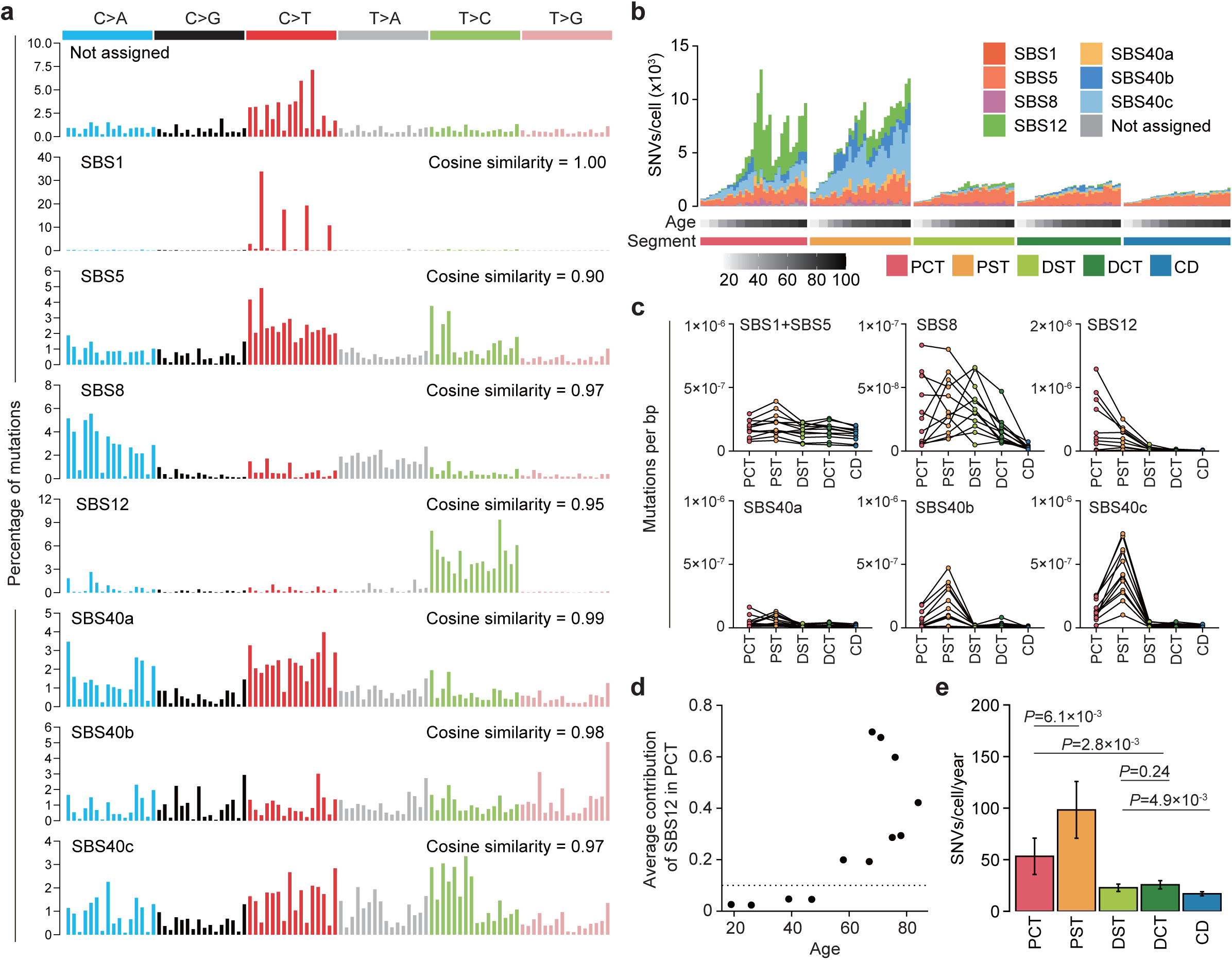
Mutational signatures in normal nephrons. a, HDP-decomposed mutational signatures assigned to corresponding COSMIC SBS signatures with cosine similarity. **b,** Landscape of mutational signatures showing absolute contributions of indicated SBS signatures in each nephron segment. Donor age is shown by a color gradient. **c,** Mutation burdens attributed to each SBS signature across nephron segments; measurements from the same donor are connected by lines. **d,** Average contribution of SBS12 in PCT for each donor, where the threshold for SBS12 positivity is set to 0.1. **e,** Mean numbers (±95% confidence interval) of SNVs/cell/year (excluding SBS12 contribution). *P* values were calculated based on the interaction term of the linear regression model.

The mutational burden attributable to age-related SBS5^28^ was largely similar across all nephron segments, increased linearly with age, and accounted for most of the mutational burden in distal tubules and CDs with additional small contributions from SBS1 and SBS8 (Fig. 2b-c). By contrast, SBS40b, SBS40c, and SBS12 were virtually confined to proximal tubules, explaining most of the mutational burdens therein (Fig. 2c). Among these, SBS40b and SBS40c, which were initially identified in ccRCC, represented major mutational signatures in proximal tubules next to SBS40a with a predominance of SBS40b (∼20%) over SBS40c (∼3%)^26^, supporting the proximal tubule origin of ccRCC proposed based on gene expression profiling^21^. In normal nephrons, SBS40a mutations showed only marginal contributions to their mutational profiles and SBS40c exhibited much larger mutational burdens than SBS40b in both PCT and PST, suggesting that they are acquired during carcinogenesis (Fig. 2c, ED_Fig. 3a-b).

**Figure 3.**
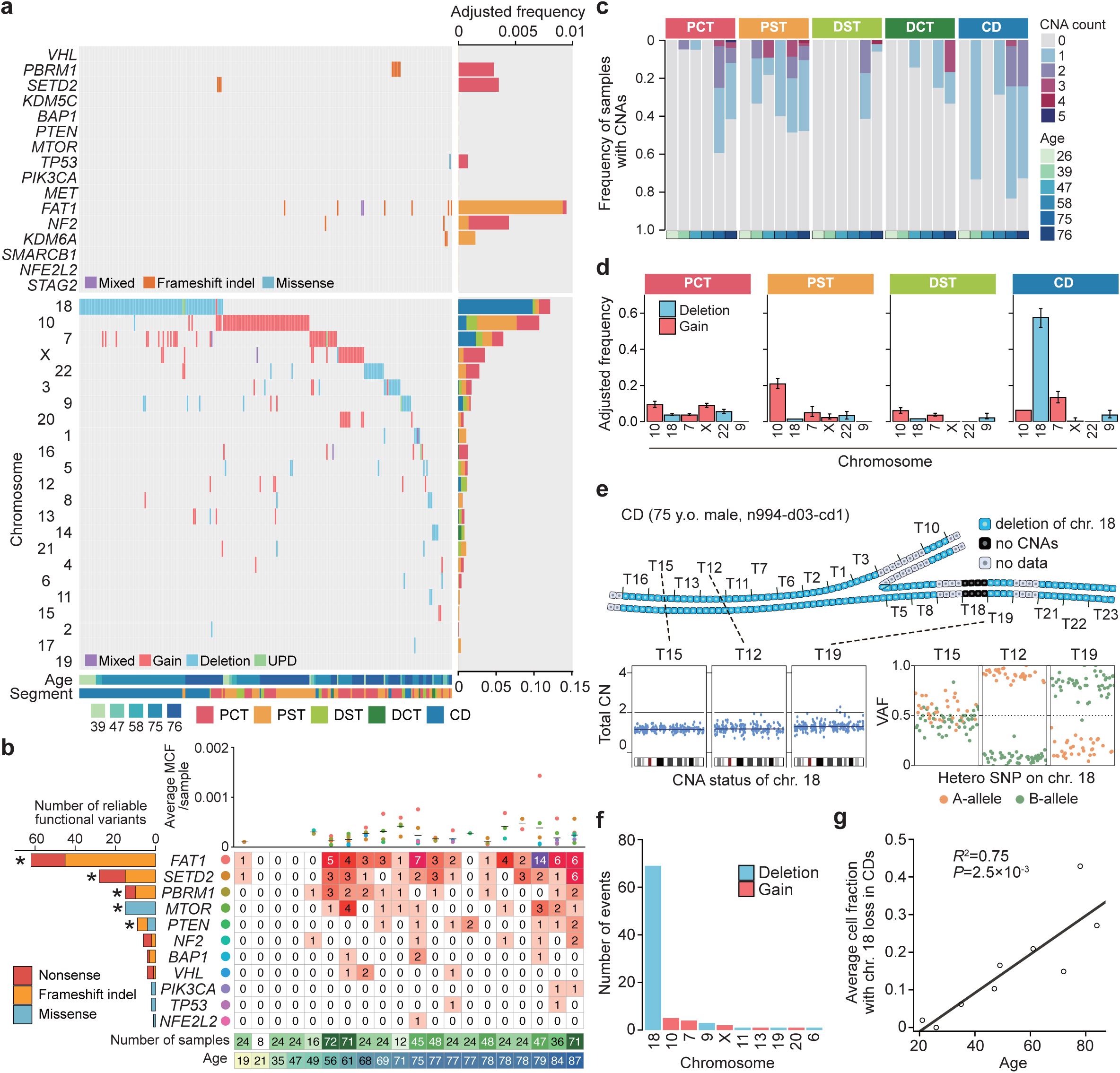
Positive selections in normal nephrons. a, Oncoplot of known RCC drivers and CNAs detected by WES in 246 of 683 LCM samples from six donors. For drivers, only highly reliable functional variants are shown. Bar plots on the right show alteration frequencies adjusted for contiguous sampling across nephron segments **(Methods)**. **b,** Number of highly reliable functional variants detected by duplex-seq of 13 candidate genes in 690 bulk samples from cortex (n=463) and medulla (n=227) (left), and a heatmap showing their distribution across donors (right). Genes under positive selection (dN/dS>1, *q*<0.1) are marked with asterisks. The upper panel shows the average mutant cell fraction (MCF) per gene for each patient; black bars indicate the mean across all detected genes. **c**, Fractions of samples carrying the indicated numbers of CNAs across segments in six donors. **d**, Frequencies of recurrent CNAs adjusted for contiguous sampling in four nephron segments. **e,** Multiple distinct clones carrying chromosome 18 loss detected along a long CD segment in a representative case (top). Bottom panels show total and allele-specific copy-number profiles labeled as A- and B-allele frequencies at heterozygous SNPs on chromosome 18 in three samples. **f,** Number of samples carrying indicated CNAs in an additional 203 CD samples newly collected from nine donors (16–24 samples/donor). **g,** Mean cell fractions carrying chromosome 18 loss in CD samples are plotted against donor age in nine donors, with a fitted regression line. *R*^2^ and *P* values were obtained using a linear regression model.

SBS12 is another mutational signature highly enriched in proximal tubules. It was also reported in ccRCC but, unlike SBS40b and SBS40c, observed almost exclusively in Japanese cases^26^. It was found in the majority of our Japanese cases (8/12) and contributed to 2.3-78.9% of the total mutational burdens in proximal tubules, which was significantly larger in PCTs than in PSTs (p=0.016) (Fig. 2d, ED_Fig. 3c-d). SBS12 was rarely detected in younger donors <40 years of age and exhibited marked inter-individual variability, accounting for a substantial fraction of the variability in mutational burden across proximal tubules (Fig. 1c and 2d). Accordingly, exclusion of SBS12 markedly improved the linearity of age-related mutation accumulation in proximal tubules and accentuated the higher baseline mutational burden in PST (Fig. 2e, ED_Fig. 3e, STable 4).

Although the mutagenic processes underlying these proximal tubule-specific signatures are largely unknown, the moderate (SBS40b and SBS40c) to strong (SBS12) transcriptional strand bias associated with their high mutational burdens suggest a contribution from extrinsic mutagens to which proximal tubules are exposed at high concentrations. Indeed, SBS40b has been linked to several candidate mutagens implicated in reduced kidney function^26^. Moreover, the high frequency and specificity of SBS12 in older Japanese donors raise the possibility that population-specific exposure to mutagens, including those present in traditional Japanese foods may contribute to the development of this signature.

We identified seven indel signatures showing high cosine similarity to COSMIC ID signatures, including ID1, ID3, ID5, ID8, ID9, ID11, and ID21 (ED_Fig. 4a). Among these, ID21, ID11, and ID1 showed largely similar mutational burdens across nephron segments albeit with slight increases in proximal tubules. By contrast, ID5, ID3, ID8 and ID9 were more abundant in proximal tubules, which are characterized by a particularly large contribution from ID5 (ED_Fig. 4b-c). ID5 is a common ID signature in ccRCC^26^, the etiology of which is still unknown.

**Figure 4.**
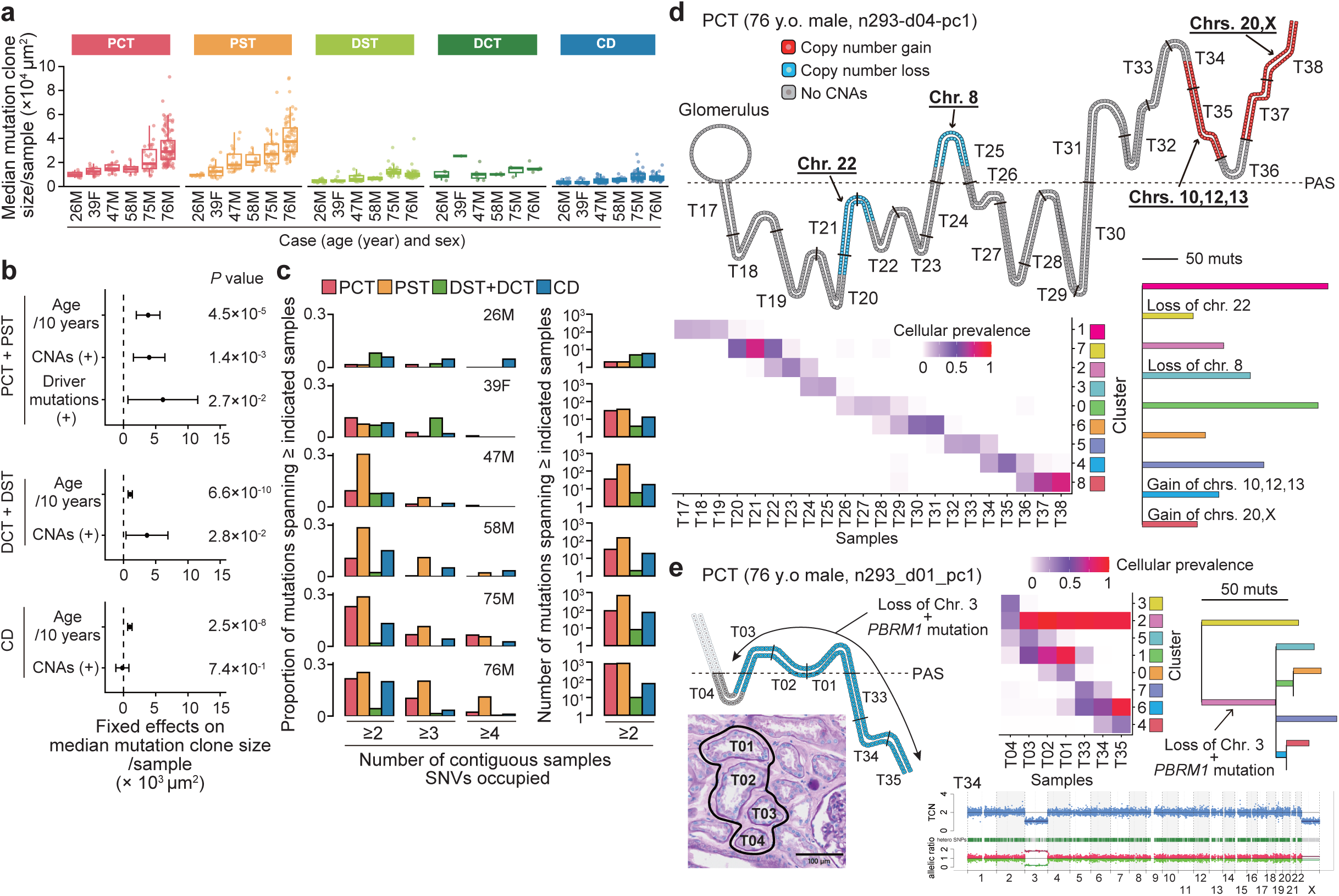
Clonal expansion in normal nephrons. a, Median mutation clone size of 683 LCM samples across five nephron segments for each donor. Mutation clone size is calculated as the mutant cell fraction (MCF) multiplied by the LCM dissection area. Box plots indicate the median (centre line), interquartile range (box) and 1.5 × IQR (whiskers). **b,** Estimated fixed effects on median mutation clone size and *P* values calculated using generalized linear mixed models (GLMM). Samples were grouped into proximal (PCT + PST), distal (DCT + DST), and collecting duct (CD) segments. Explanatory variables included age (per 10 years), presence of CNAs, and presence of driver mutations (highly reliable functional variants in RCC driver genes). Error bars indicate 95% confidence intervals. **c,** Proportion (left) and number (right) of SNVs and indels detected in ≥ a given number of contiguous samples in indicated nephron segments. **d,** Representative examples of a set of contiguously collected LCM samples from a long PCT segment. Samples carrying CNAs are indicated by colors corresponding to the affected chromosomes (top). For each sample, the cellular prevalence of clones inferred by PyClone-VI is shown as a color gradient (bottom left), together with their inferred phylogenetic relationships (bottom right). **e,** Representative examples of extensive clonal expansion in aged nephrons, showing a schematic nephron architecture, PAS staining (at an indicated level), and total (blue) and allele-specific (red and green) copy number profiles for the indicated samples. Also shown is the cellular prevalence of mutational clusters inferred by PyClone-VI, along with corresponding phylogenetic trees.

### Positive selections in normal nephrons

We next evaluated positive selection in distinct nephron segments using WES of an additional set of LCM samples independently collected for this purpose (Fig. 3a). In total, we obtained 683 LCM samples spanning five distinct nephron segments from six donors representing different age groups. Of these, 500 were organized into 100 sample sets, each consisting of 2-22 contiguous samples (median=4) (ED_Fig. 5a), while the remaining 183 were collected as non-contiguous, solitary samples (ED_Fig. 5b-c). The number of tissue slices used for microdissection was adjusted so that each LCM sample contained comparable numbers of epithelial cells (n=∼200-300) (Methods for details and SFig. 1a-b).

**Figure 5.**
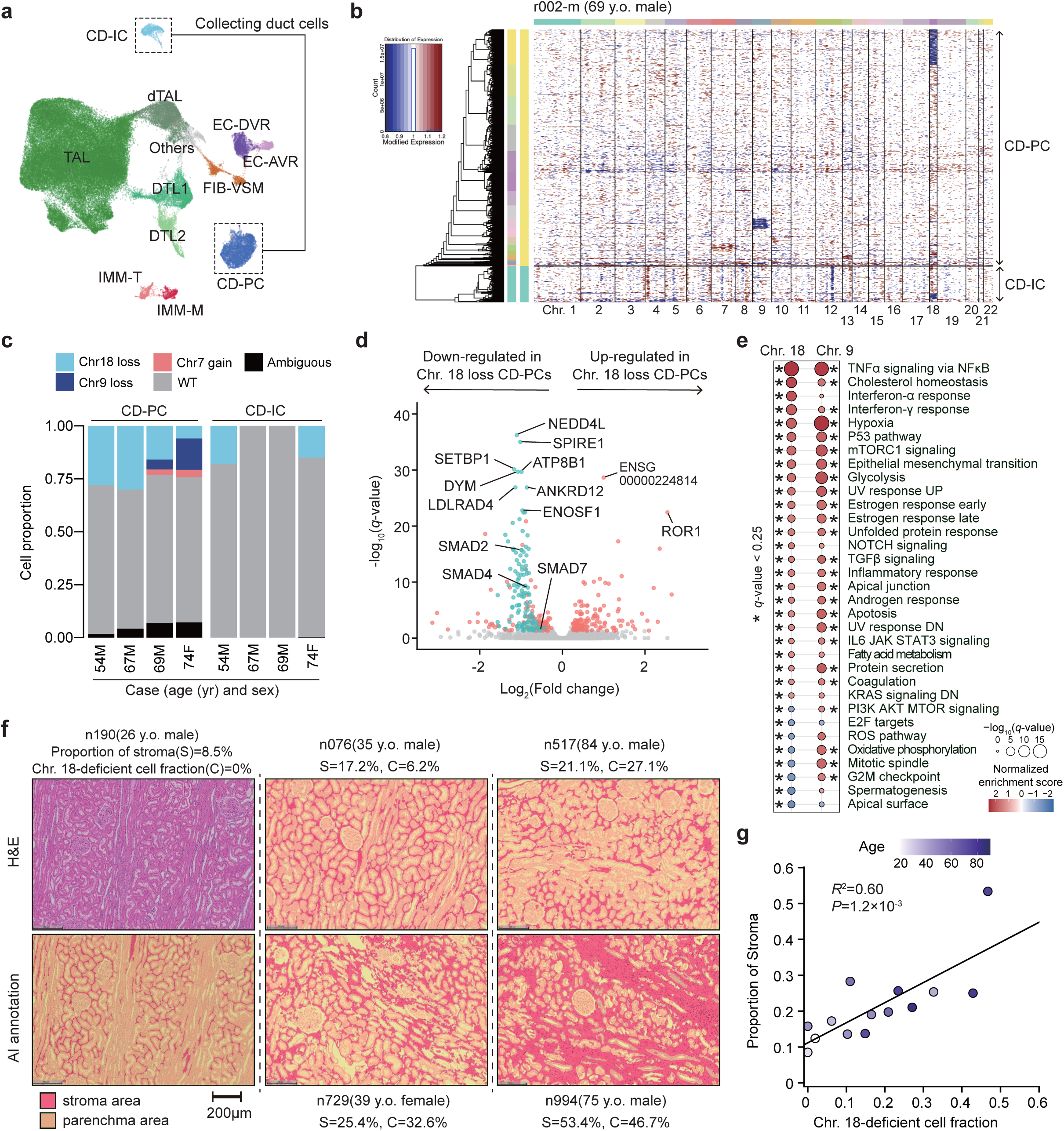
Impact of CNAs on gene expression in CDs. **a,** UMAP combining snRNA-seq data from four donors. CD-PC, principal cell of the collecting duct; CD-IC, intercalated cell of the collecting duct; DTL1 and DTL2, descending thin-limb cell types 1 and 2; TAL, thick ascending limb cell; dTAL, degenerative thick ascending limb cell; IMM-T, T cell; IMM-M, macrophage; FIB-VSM, fibroblast and vascular smooth-muscle cell; EC-DVR and EC-AVR, endothelial cells of descending and ascending vasa recta. **b,** A representative result of snRNA-seq-based copy number inference for a medullary sample from a 69-year-old male donor, delineating distinct clusters of CD-PC carrying chromosome 18 loss showing global reductions in expression of chromosome 18 genes, as well as those carrying chromosome 9 loss and chromosome 7 gain in CD-PC. **c,** Inferred CNAs and their cell fractions in CD-PC and CD-IC based on gene expression profiles. Clusters comprising fewer than 2.5% of total CD-PC and CD-IC were collectively categorized as ambiguous. **d,** Volcano plot of DEGs identified in pseudo-bulk analysis comparing CD-PCs with and without chromosome 18 loss. Significant DEGs (*q*<0.05 and |log2FoldChange|>0.25) are colored red or green, with genes on chromosome 18 highlighted in green. **e,** Gene set enrichment analyses of MSigDB Hallmark gene sets comparing CD-PCs with and without chromosome 9 or 18 loss. Significant gene sets (*q*<0.25) are marked with asterisks. Only pathways significantly enriched in the chromosome 18-loss clones are shown. **f**, Representative AI-assisted histological images. The left column displays H&E staining (top) and corresponding AI annotation (bottom) of a 26-year-old male, highlighting the minimal presence of stroma. The middle and right columns show representative younger and older cases, respectively, illustrating the spectrum of stromal proportion (S) and chromosome 18-deficient cell fraction (C). **g,** Proportions of stroma are plotted against chromosome 18-deficient cell fractions, with a fitted regression line. *R*^2^ and *P* values were obtained using a linear regression model.

WES was performed with a median depth of 51.2×. After excluding redundant variants detected in contiguous samples, we identified a total of 24,433 SNVs and 2,385 indels as independent variants in 683 LCM samples, which were then used to investigate driver mutations. Based on dN/dS analysis, no genes were shown to be significantly positively selected either in genome-wide testing or when restricting the analysis to 16 known RCC driver genes^22,29,30^ (STable 5). A small number of nonsynonymous mutations (n=59, 0.22%) were detected within these RCC driver genes in the minority (9.4%) of samples. However, only 13 of these mutations represented highly reliable functional variants registered in the COSMIC database (Methods for details and STable 6), including nonsense or frameshift mutations and known recurrent hotspot mutations, and the driver role of the remaining mutations was unclear due to their rarity and the lack of evidence for positive selection (dN/dS not significantly greater than 1.0) (Fig. 3a, STable 5). In particular, *VHL* mutations, which account for the vast majority of ccRCC (>80%)^31^, were not detected in our WES dataset. To further investigate RCC-related mutations in normal nephrons at higher sensitivity, we performed deep duplex sequencing targeting 13 common RCC driver genes^22,30–32^ (STable 7; Methods). In total, we analyzed 690 bulk samples (mean volume 0.2 mm^3^) taken from cortex (n=463) and medulla (n=227) in 20 donors with a mean duplex depth of 6,119× (ED_Fig. 5d-g, SFig. 2a). We identified 647 non-synonymous mutations and 104 indels across the 13 genes (ED_Fig. 5h, STable 8), of which five showed significant positive selection (dN/dS>1) (STable 5). However, except for truncating and hotspot variants, dN/dS values for the vast majority of mutations only marginally exceeded 1.0, suggesting that a minority of these mutations were functional. In line with this, only 148 of all non-synonymous variants (n=751) represented highly reliable functional variants (Fig. 3b, STable 8). These results suggest that the expansion of RCC-related driver mutations beyond the threshold of duplex sequencing (∼120 cells) is very rare in normal kidneys, but nevertheless, showed a significant enrichment in cortex samples (odds ratio (OR)=3.3, 95% confidence interval (CI): 1.9-5.9), supporting the proximal tubule origin of major RCCs.

CNAs were also investigated based on the WES data (Methods). CNAs were quite common, detected in more than one third (240/683) of LCM samples, and consisted predominantly of arm-level alterations (Fig. 3a, SFig. 3a and STable 9). CNAs, particularly multiple CNA lesions, were strongly correlated with donor age (p=1.5×10^-3^) and were highly enriched in aged donors (Fig. 3c). CNAs most frequently affected chromosomes 18, 10, and 7, followed by chromosomes X, 22, 3, and 9, where the patterns of gains (chromosomes 10, 7, and X) and losses (chromosomes 18, 22, and 9) were largely consistent for each chromosome (Fig. 3a), supporting their roles in positive selection. These CNAs were rarely observed in RCC, except for losses of chromosome 3 (ccRCC) and losses of chromosome 9, as well as gains of chromosome 7 (ccRCC and pRCC). Chromosome 3p losses, which are hallmarks of ccRCC, were rarely observed in normal nephrons.

Most CNAs were enriched in proximal tubules, except for chromosome 18 loss and chromosome 7 gains. Chromosome 7 gain was observed across multiple segments, whereas chromosome 18 loss was virtually confined to CDs (Fig. 3a,d). Detected in up to 80% of LCM samples in aged donors, chromosome 18 loss virtually explained most CNA events in CDs (Fig. 3a,c). A representative example was observed in a 75-year-old male, in whom chromosome 18 losses were observed in 17 out of 18 contiguous CD samples, typically at high cell fractions (Fig. 3e). Remarkably, the parental origin of the retained allele (A or B) in each LCM sample alternated along this CD segment spanning 20 slides. Moreover, marked discrepancies between total and allele-specific copy numbers were observed in T15 and also in samples from different donors (ED_Fig. 6a-c). These findings suggest extreme parallel evolution of numerous independent clones lacking chromosome 18 in CDs, even within such small samples comprising ∼200-300 cells.

To further assess the frequency and age-dependence of chromosome 18 loss in CDs, we collected an additional set of LCM samples (n=203) from nine donors (16–24 samples/donor) from different age groups, followed by WES (Fig. 1b, STable 1). We confirmed that chromosome 18 loss was the most frequent CNA in CDs in aged donors (Fig. 3f-g). The estimated cell fractions carrying chromosome 18 loss linearly increased with age, replacing approximately 30% of CD cells by 80 years of age (Fig. 3g), which was substantially higher than the rough estimate of the frequency of proximal tubules carrying chromosome 10 gain (ED_Fig. 6d). Taken together, these results indicate that cells carrying chromosome 18 loss are strongly positively selected in CDs and progressively replace surrounding wild-type CD cells over years, ultimately contributing to age-related remodeling of nephron architecture in the elderly.

### Clonal expansion in aged nephrons

Clonal expansion in normal nephrons was assessed by variant allele frequencies (VAFs) of mutations detected in WES. To adjust the effect of sample size, we calculated ‘mutation clone size’ for each mutation by multiplying the mutated cell fraction (=2×VAF) and corresponding sample size (Fig. 4a). In the youngest donor (26 years), median mutation clone size in each sample was uniformly low across all segments, only slightly above the detection threshold (for VAF≥0.08) and mostly well below 0.25 with no evidence of monoclonal expansion. By contrast, in aged donors, proximal tubule samples exhibited markedly increased mutation clone sizes, together with mutation numbers, with many showing monoclonal expansion (Fig. 4a, ED_Fig. 7a-b, SFig. 4). On average, PSTs in the aged donor (76 years) exhibited 4.04-fold (95% CI, 3.38–4.77) increase in median mutation clone size relative to that in the 26-year-old donor, which was larger than the size in PCTs (2.99 times; 95% CI, 2.42–3.78). Notably, virtually all mutations in proximal tubule samples from aged donors exhibited larger mutation clone size than that in the youngest donor (Fig. 4a). In distal tubules and CDs, in contrast, the age-related increase in mutation clone size remained substantially smaller, compared with that in proximal tubules (Fig. 4a). In fact, in multivariable analysis based on the Generalized Linear Mixed Model (GLMM), donor age was a significant predictor of mutation clone size measured by median mutation sizes across all segments, but the effect was much larger in proximal tubules than in other segments (Fig. 4b and STable 10). CNAs also significantly affected median mutation clone sizes in proximal and distal tubules, but not in CDs. In proximal tubules, the cumulative effect of advanced age (75–76 years) on median mutation clone size was much larger than that of CNAs, although the effects of age and CNAs were comparable in distal nephrons. Almost exclusively detected in proximal tubules, highly reliable driver mutations were also significantly associated with mutation clone size, but the estimated effect size remained similar to that of CNAs (Fig. 4b).

Clonal expansion was further assessed by identifying mutations shared between adjacent samples using 100 sets of contiguously collected samples (median, 4; range, 2–22), comprising a total of 500 samples (Fig. 4c, ED_Fig. 5a-c). In the youngest donor, the number of shared mutations was uniformly low across different segments, where small numbers of shared mutations across contiguous samples likely represent the mosaicism established during early nephrogenesis rather than clonal expansion thereafter (Fig. 4c). However, in aged donors (75 and 76 years), mutations in proximal tubules, particularly those in PSTs, were more frequently shared across multiple contiguous samples, compared to those in other segments (Fig. 4c). In a PCT sample from the 76-year-old donor, as many as nine distinct clones, including those carrying different CNAs, expanded over ≥ four contiguous samples, packed side by side, intermingling with adjacent clones (Fig. 4d). By contrast, in distal tubules, the frequency of shared mutations, particularly those spanning ≥3 samples, was almost consistently lower than that in proximal tubules (6.8–20.3%) in aged donors (Fig. 4c). CDs showed an intermediate pattern of mutation sharing: the frequency of mutations shared between two adjacent samples was comparable to that in proximal tubules and much higher than that in distal tubules. Mutations were mostly confined to two samples even in aged donors, but a small fraction of mutations extended to ≥3 contiguous samples, indicating limited but appreciable expansion of clones in CDs (Fig. 4c). Combining mutations detected in multiple contiguous samples, we identified three clones in proximal tubules showing particularly large expansion in two aged donors (Fig. 4e, ED_Fig. 7c-e). Carrying one or more CNAs, all these clones extended spanning up to seven contiguous samples, giving rise to multiple subclones with or without additional CNAs (Fig. 4e, ED_Fig. 7d-e). Among these, of particular interest is the clone found in a PCT. Harboring chromosome 3 loss and a *PBRM1* frameshift mutation, a characteristic combination commonly seen in ccRCC, this is thought to represent an early lesion predisposed to ccRCC, although no morphological changes were observed.

### Impact of CNAs on gene expression in CDs

Finally, we investigated the transcriptional consequences of recurrent CNAs in collecting ducts (CDs), focusing on chromosome 18 loss. We performed single-nucleus RNA sequencing (snRNA-seq) on fresh-frozen medullary tissues obtained from four patients with ccRCC aged 54–74 years (Fig. 5, ED_Fig. 8– 10, SFig. 5; Methods). After quality control, 81,614 nuclei were analyzed and grouped into 12 clusters representing distinct renal cell populations (Fig. 5a, ED_Fig. 8a-c). CD principal cells (CD-PCs) and intercalated cells (CD-ICs) accounted for 2.9–11.9% and 0.4–1.4% of medullary cells, respectively. Using expression-based copy number inference (inferCNV^33^), we identified distinct clusters of chromosome 18-deficient cells within the CD-PC compartment in all four samples and within the CD-IC compartment in two samples (Fig. 5b-c, ED_Fig. 9a). Additional clusters associated with chromosome 9 loss and chromosome 7 gain were also detected in CD-PCs from two older individuals.

To evaluate the transcriptional effects of these CNAs, we performed pseudo-bulk differential expression analyses comparing CD-PCs with and without chromosomal abnormalities. Due to the limited number of CNA-positive cells, subsequent analyses focused on chromosome 18 and chromosome 9 losses. For both lesions, the most significantly downregulated genes were highly enriched on the affected chromosome, accounting for 78 and 67 of the top 100 downregulated genes for chromosome 18 and chromosome 9 losses, respectively (Fig. 5d, ED_Fig. 10a, STable 11), confirming the strong allele-dosage effect of chromosomal loss on gene expression that underlies the gene expression-based copy number inference. Among these, several genes on chromosome 18 have been implicated in renal homeostasis and tissue injury responses. For example, *NEDD4L,* which was most significantly downregulated in chromosome 18-deficient cells, encodes an E3 ubiquitin ligase involved in the negative regulation of the epithelial sodium channel (ENaC), a key regulator of sodium and fluid balance expressed in CD-PCs. Consistent with this, genetic inactivation of the murine orthologue *Nedd4-2* results in increased ENaC activity, sodium retention, and salt-sensitive hypertension^34–36^. Other genes on chromosome 18, including *SMAD2*, *SMAD4*, *SMAD7*^37–40^ and also *NEDD4L*^34,41,42^, participate in TGF-β/Smad signaling implicated in tissue repair, inflammation, and fibrosis^37^ (Fig. 5d).

To assess the broader functional consequences of chromosome 18 loss, we performed gene set enrichment analysis (GSEA) of pseudobulk samples combining all CD-PCs. Compared with chromosome 18-proficient CD-PCs, chromosome 18-deficient cells exhibited enrichment of multiple inflammatory and profibrotic programs, including TNF-α signaling via NF-κB, interferon-α and interferon-γ responses, IL6–JAK–STAT signaling, TGF-β signaling, epithelial–mesenchymal transition (EMT), and inflammatory response pathways (Fig. 5e, ED_Fig. 10b). Similar transcriptional changes were also observed in CD-PCs carrying chromosome 9 loss (Fig. 5e), with substantial overlap between the pathways enriched in the two CNA categories. These shared signatures included TNF-α signaling via NF-κB, interferon responses, hypoxia, TP53 pathway, mTORC1 signaling, EMT, glycolysis, inflammatory response, TGF-β signaling, apoptosis, autophagy, and IL6–JAK–STAT signaling, suggesting a common adaptive and/or selective program associated with losses of these chromosomes in aging CDs. Notably, many of these pathways overlap with transcriptional programs reported in the unique subset of proximal and distal tubule cells induced after acute kidney injury^43–45^. Given this overlap in transcriptional programs, we next examined whether accumulation of chromosome 18-deficient cells was associated with tissue remodeling in vivo. In healthy young donors, the interstitial space between renal tubules is inconspicuous but gradually expands with aging as fibrosis accompanied by chronic inflammation progresses (Fig. 5f). We therefore quantified the interstitial area as a surrogate measure of fibrosis and compared it with the fraction of chromosome 18-deficient cells. Although interstitial expansion generally increased with age, the extent of fibrosis correlated significantly with the fraction of chromosome 18-deficient cells irrespective of age (*R*² = 0.60, *p* = 1.2 × 10⁻^2^) (Fig. 5g, ED_Fig. 10c-d). Together, these findings suggest that recurrent chromosome 18 loss in CD cells is associated with coordinated transcriptional changes involving inflammatory and profibrotic pathways and may contribute to age-associated kidney fibrosis.

## Discussion

Through extensive sequencing analyses of a large number of microdissected samples, we have delineated how age-associated somatic mosaicism is established within spatially constrained and compartmentalized tissues like kidney nephrons. In contrast to somatic mosaicism in other non-cancerous tissues, in which cancer-related mutations often become pervasive, RCC-related mutations were only infrequently detected even in aged normal nephrons and largely confined to proximal tubules. By contrast, CNAs are quite common and appear to play a larger role in positive selection than has been reported in other tissues. However, with the exception of losses of chromosome 3 and 9, they are rarely observed in RCCs. Thus, their roles in renal carcinogenesis or aged kidney functions are unclear and so is the mechanism of the clonal selection of these segment-specific CNAs.

In nephrons, a relatively small population of cells is imprisoned in spatially fixed tubular compartments^46^. As a result, their population heterogeneity is expected to decrease with aging by repeated neutral drifts with accelerated cell turnover (Moran model), giving rise to progressively larger clones at advanced age^47,48^. CNAs and driver mutations also contribute to the increased clone size (Fig. 4b). However, their effects were much smaller than that of donor age, accounting for only 13.9–21.3% of the age effect in proximal tubules at 75–76 years of age, suggesting that clone size in nephrons is influenced more by cell turnover rate than driver events. According to this model, renal tubules, particularly proximal tubules, may undergo more active cell turnover during their lifetime than previously estimated from Ki67 staining and BrdU clearance. The increased cell turnover in proximal tubules may reflect their higher sensitivity to hypoxia and larger exposures to harmful substances and is also supported by very high mutational burdens and the accelerated cell proliferation of proximal tubule cells after acute kidney injury^46,49,50^. By contrast, likely due to reflecting their slow cell turnover, CD cells showed a much smaller increase in their clone size, even though a substantial fraction of CD cells were replaced by numerous clones with strong fitness advantage conferred by chromosome 18 loss.

Finally, one of the most striking observations was the remarkable parallel evolution of chromosome 18-loss clones in collecting ducts. The exact mechanisms underlying both the recurrent selection and expansion of chromosome 18 loss clones could not be directly addressed in the current study and therefore remain largely speculative. Nevertheless, several observations provide clues to these questions. Chromosome 18- and chromosome 9-deficient cells exhibited enhanced responses to TGF-β and multiple pro-inflammatory cytokines, together with activation of transcriptional programs characteristic of injured and aging kidneys. These findings suggest that heightened responsiveness to the chronic inflammatory and profibrotic milieu of the aging kidney may contribute to the selective advantage of chromosome-deficient cells. At the same time, chromosome 18 loss was associated with reduced expression of several genes implicated in TGF-β/Smad signaling, including *NEDD4L, SMAD2, SMAD4,* and *SMAD7*, and the burden of chromosome 18-loss cells correlated with the extent of tubulointerstitial fibrosis. In particular, NEDD4L occupies a notable position at the intersection of collecting duct sodium handling and TGF-β/Smad signaling, suggesting that chromosome 18 loss may have consequences for both epithelial homeostasis and renal tissue remodeling. Although the direction of causality remains unresolved, our findings point to a previously unrecognized interplay between age-associated clonal selection, renal aging, and tissue remodeling.

## Methods

### Study design

No statistical methods were used to predetermine sample size since this is an exploratory study. The experiments were not randomized and, unless otherwise stated, the investigators were not blinded to allocation during experiments and outcome assessment. The annotation of nephron segments and position based on H&E and PAS staining was conducted prior to immunohistochemistry and the analysis of sequencing results.

### Participants and materials

We enrolled a total of 32 donors from three institutions between July 22, 2020, and August 1, 2025: Kyushu University Hospital, Shiga University of Medical Science Hospital, and Iwate Medical University Hospital. Specifically, we obtained 29 surgical specimens from patients at Kyushu University Hospital who underwent surgery for renal tumors. Iwate Medical University Hospital and Shiga University of Medical Science Hospital provided autopsy specimens, comprising 2 and 1 donors, respectively. Donor characteristics are summarized in STable 1. This study was reviewed and approved by the Institutional Ethics Committee at all participating institutions. Written informed consent was obtained from all donors or their families. Normal kidney tissue was collected, either fixed with PAXgene Tissue FIX (QIAGEN, cat. 765312) or snap-frozen. Paired peripheral blood samples were also collected. For autopsy specimens where peripheral blood was unavailable, the spleen was collected and fixed with PAXgene Tissue FIX instead. In patients with renal tumors, normal tissues were collected from the surgical margins, and the resection surfaces were pathologically confirmed to be negative for tumor cells.

### Immunohistochemistry to validate accuracy of nephron segment annotation

Nephron segments were discriminated based on their staining properties with H&E and PAS (three types: proximal tubule, distal tubule, and collecting duct) and their morphology (two types: convoluted and straight parts). To evaluate the accuracy of H&E/PAS-based classification, we performed immunohistochemistry (IHC) using segment-specific antibodies and compared the results with the H&E/PAS-derived annotations (ED_Fig. 1a-c). Serial sections were prepared from a PAXgene-fixed normal kidney block, and IHC was performed on sections consecutive to those used for H&E and PAS staining. For IHC, sections were stained using the ImmPRESS® HRP Horse Anti-Mouse IgG Polymer Detection Kit, Peroxidase (Vector Laboratories) or the ImmPRESS® HRP Horse Anti-Rabbit IgG Polymer Detection Kit, Peroxidase (Vector Laboratories), following the manufacturer’s protocol. Primary antibodies against UMOD (Atlas Antibodies, cat. HPA043420, 1:300 dilution), SLC12A1 (Atlas Antibodies, cat. HPA018107, 1:600 dilution), SLC12A3 (Atlas Antibodies, cat. HPA028748, 1:200 dilution), AQP2 (Atlas Antibodies, cat. HPA046834, 1:200 dilution), CD10 (Thermo Fisher Scientific, cat. MA514050, 1:200 dilution), and CD34 (Abcam, cat. ab81289, 1:2500 dilution) were used. For antigen retrieval, sections for CD10, UMOD, SLC12A1, SLC12A3, and AQP2 antibodies were incubated in Antigen Unmasking Solution, Citric Acid Based (Vector Laboratories) at 55 °C for 30 minutes, but no retrieval was performed for the CD34 antibody. Sections were incubated with primary antibodies at 4 °C overnight. Prior to IHC evaluation, each nephron was classified based on H&E and PAS staining patterns. Subsequently, for nephrons in serial sections where all markers could be evaluated, marker-based validation was conducted. To quantify the consistency between the staining properties and the molecular profiles, we calculated a concordance rate for each segment. Concordance was assessed as follows: “True” if only the segment-specific marker was expressed; “Ambiguous” if both the target marker and non-target markers were co-expressed, or if no markers were detected; and “False” if only one or more non-target markers were expressed. The concordance rate was defined as the ratio of “True” samples to the total number of evaluated samples. Based on this definition, the concordance rates in a single case were 100% (47/47) for the proximal tubule, 98% (41/42) for the distal tubule, and 96% (23/24) for the collecting duct. (ED_Fig. 1d).

### Preparation of tissue sections and LCM

PAXgene-fixed samples were embedded in paraffin, ensuring that the sagittal section face was exposed to facilitate the discrimination between convoluted and straight segments based on their spatial orientation. PAXgene-fixed, paraffin-embedded (PFPE) samples were contiguously sectioned at a thickness of 8 μm and mounted on frame slides with POL membranes (Leica). Slides for H&E and PAS staining were prepared approximately every 20 sections. While a minimum of 20 contiguous sections, excluding those used for H&E and PAS staining, were required to reliably identify convoluted and straight segments, 40 consecutive sections were typically prepared to ensure a robust sample set. PFPE slides were deparaffinized and stained with Hematoxylin Solution, Gill No. 2 (Sigma-Aldrich) for 10 seconds, followed by two 10-second washes with distilled water. The slides were then incubated with 1% Eosin Y solution (FUJIFILM, 1:6 dilution) for 5 seconds, followed by a 10-second wash with distilled water, two 5-second washes with 70% ethanol, and two 5-second washes with 100% ethanol. Slides were imaged using the BZ-X800 (Keyence) or VS200 (Evident). LCM was performed using the LMD7000 or LMD7 system (Leica).

### Quantitative input control for LCM samples in WES

Cell density varies across nephron segments, necessitating adjustments to the dissection area to maintain a target of 200–300 cells per sample (SFig. 1a). Since library yield reflects the input DNA amount under identical library preparation conditions, we monitored these yields to ensure they fell within a consistent range. Empirically, an input of approximately 200 cells yielded roughly 500–700 ng of DNA library. In practice, library yields for the majority of samples consistently fell within the 500– 1,000 ng range (SFig. 1b).

### Preparation of punch biopsies for duplex sequencing

PAXgene-fixed samples were sagittally sliced to an average thickness of 500 μm using the Thomas Stadie-Riggs Tissue Slicer (Thomas Scientific). The sliced tissues were then punched out using a 1-mm biopsy punch (Kai Industries) and further bisected with a scalpel (ED_Fig. 5d). The punched regions were confirmed to be free of cancer or dysplasia by evaluating adjacent H&E-stained sections (SFig. 2). Regarding the distinction between the cortex and medulla, the punched regions containing renal labyrinths were annotated as the cortex.

### Cell lysis and DNA extraction

For LCM samples, cell lysis was performed as previously described by Ellis et al.^51^, using an in-house buffer followed by DNA capture on magnetic beads and ethanol washing. The beads were then resuspended in low-TE buffer, and the suspension was directly subjected to library construction without DNA quantification. For punch biopsy samples intended for duplex sequencing, the protease concentration in the in-house buffer was increased 10-fold, and the incubation at 50 °C was extended to 60 minutes. Two rounds of magnetic bead purification were performed, and the DNA was eluted in low-TE buffer. DNA from peripheral blood and spleen samples was extracted using the Gentra Puregene Kit (QIAGEN) according to the manufacturer’s instructions. The concentration of the purified DNA was quantified using the Qubit dsDNA BR Assay Kit (Thermo Fisher Scientific).

### Library construction for NanoSeq, WES and duplex sequencing

For NanoSeq, DNA library construction was adapted from the original protocol^24^. Specifically, A-tailed DNA fragments were ligated to UMI-containing adapters using the Twist cfDNA Library Preparation Kit (Twist).

For WES and duplex sequencing of LCM and biopsy samples, library construction began with DNA fragmentation using the xGen DNA Library Prep EZ Kit (IDT), with the fragmentation reaction arrested at 32 °C. The fragmented DNA was purified by adding a 2.5× volume of AMPure XP beads (Beckman Coulter), followed by two washes with 80% ethanol, and then resuspended in low-TE buffer. Subsequently, strand-specific 8-bp UMI adapters were ligated using the xGen cfDNA & FFPE DNA Library Preparation Kit (IDT). For matched control DNA from peripheral blood and spleen, DNA libraries were prepared using either the xGen DNA Library Prep EZ Kit (IDT) or by fragmenting the DNA with a Focused-ultrasonicator E220 (Covaris) followed by the xGen cfDNA & FFPE DNA Library Preparation Kit (IDT), according to the manufacturer’s instructions. NanoSeq libraries were quantified by qPCR using the KAPA Library Quantification Kit (KAPA Biosystems), whereas libraries for WES and duplex sequencing were assessed using the D1000 ScreenTape (Agilent).

### NanoSeq

Sequencing was performed on either the DNBSEQ-G400RS or the DNBSEQ-T7 (MGI Tech) in 150-bp paired-end mode. Data processing was conducted using the NanoSeq pipeline (v.3.5.4) (https://github.com/cancerit/NanoSeq), with two modifications to the default settings. First, the UMI length was increased from 3 bp to 5 bp. Second, the SNP and noise masks were optimized to exclude SNPs and noise specific to the Japanese population. These masks were generated using the ToMMo 61KJPN-SNV/INDEL Allele Frequency Panel (v20250616)^52^, applying filtering criteria equivalent to the original masks. Specifically, SNP mask candidates were defined as common SNPs on autosomes and the X chromosome with an allele frequency (AF) >0.1% and a “PASS” flag, and SNPs on the Y chromosome and mitochondrial DNA with an AF >0.1%. Noise mask candidates included indel calls with an AF >1%, and SNP calls with an AF >0.1% that were not flagged as “PASS”. These candidate lists were lifted over from hg38 to hg19 and integrated into the default NanoSeq masks. The final SNP and noise masks covered 31,994,146 bp and 42,290,445 bp, respectively.

### Signature analysis

Mutational signatures were extracted using the Hierarchical Dirichlet Process (HDP) package (https://github.com/nicolaroberts/hdp). The HDP analysis was conducted with 20 independent chains running for 40,000 iterations each, with a burn-in period of 20,000 iterations. From each chain, 100 posterior samples were collected at regular intervals after the burn-in period, and clusters exhibiting a cosine similarity of 0.9 or higher were merged into a single component. For analyses incorporating prior information, a prior pseudocount of 1,000 was applied.

First, we performed signature extraction on the 96 substitution classification using the COSMIC v3.4 SBS signature catalog as prior information. This identified seven primary components (Component P5 = SBS5, Component P18 = SBS12, Component P50 = SBS40c, Component P49 = SBS40b, Component P48 = SBS40a, Component P11 = SBS8, and Component P1 = SBS1) (Fig. 2a). Since all seven components showed high cosine similarity (>0.90) to their corresponding COSMIC SBS signatures, no further deconvolution was performed. Clusters that could not be stably assigned to any main component were defined as “Not assigned” (Component 0).

Next, we performed *de novo* signature extraction on the 192 substitution classification to account for transcriptional strand bias. Five main components were stably extracted. To verify the consistency of these signatures with those identified in the 96 substitution analysis (SBS1, 5, 8, 12, and 40a–c), the 192 substitution classification was collapsed back to the 96 substitution classification. Four of the five components (Component 1 = SBS5, Component 2 = SBS40c, Component 3 = SBS12, and Component 4 = SBS40b) exhibited very high cosine similarity (>0.95) to the SBS signatures extracted from the 96 substitution analysis (ED_Fig. 2a).

Finally, for the 83 indel classification, we performed signature extraction using the COSMIC v3.4 ID signature catalog as prior information. This resulted in the extraction of six main components, all of which showed high cosine similarity (>0.95) to signatures in the COSMIC v3.4 ID signature catalog (Component 1 = ID1, Component 3 = ID3, Component 5 = ID5, Component 8 = ID8, Component 11 = ID11, and Component 21 = ID21). Similarly, Component 0 also exhibited a very high cosine similarity (0.996) and was therefore defined as ID9 (ED_Fig. 4a).

### Whole-exome sequencing

All indexed libraries underwent hybridization and capture using the xGen^TM^ Exome Hyb Panel v2 (IDT). Sequencing was performed on a DNBSEQ-G400RS (MGI Tech) in 150-bp paired-end mode. BAM file generation and mutation calling were performed using the Genomon pipeline (v.2.6.3), as previously described^53^, with a modification to GenomonMutationFilter (v.0.4.0) (https://github.com/kwatanabe-ku) to assess artifacts related to cruciform DNA formation. The average sequencing depth for all LCM samples was 57.6× (range, 18.2 to 147.1×). Called mutations were further filtered if they did not meet the following criteria: (i) variant reads ≥3; (ii) variant allele frequencies (VAFs) ≥0.08 in the sample; (iii) VAFs ≤0.02 in the paired normal control; (iv) strand ratio ≠0 or 1; (v) Fisher’s test *P* value ≤10^-1^; (vi) EBcall *P* value ≤10^-4^ using 20 whole-exome sequencing data from non-paired spleen or interstitial tissue; (vii) the ratio of cruciform-derived reads among variant reads <0.5; (viii) maximum variant reads detected in the same 20 non-paired control samples used for EBcall ≤2; (ix) variants were not located within segmental duplications documented in the GenomicSuperDups database. To further eliminate sequencing errors, mutations detected across independent nephron samples within the same individual were subjected to visual inspection on the Integrative Genomics Viewer^54^.

### Whole-genome sequencing

We selected LCM samples with a median VAF ≥ 0.25 in WES and sufficient remaining DNA library for additional whole-genome sequencing (WGS). Sequencing and BAM file generation were performed following the same protocols as described for WES. The average sequencing depth was 31.8× (range, 20.3 to 48.0×). Called mutations were further filtered if they did not meet the following criteria: (i) depth ≥8 in both the sample and the paired normal control; (ii) variant reads ≥3 in the sample and ≤1 in the paired normal control; (iii) variant allele frequencies (VAFs) ≥0.05 in the sample; (iv) strand ratio ≠0 or 1; (v) Fisher’s test *p*-value ≤10^-1^; (vi) EBcall *p*-value ≤10^-4.5^ using 20 whole-genome sequencing datasets from non-paired spleen or interstitial tissue; (vii) the ratio of cruciform-derived reads among variant reads <0.5.

### Duplex sequencing

We selected 13 genes commonly mutated in major RCC subtypes (ccRCC^31,32^, pRCC^30^, and chRCC^22^) for duplex sequencing, including *VHL*, *PBRM1*, *SETD2*, *BAP1*, *TCEB1*, *MET*, *FAT1*, *NF2*, *PTEN*, *mTOR*, *PIK3CA*, *NFE2L2*, and *TP53*. For each sample, 120 ng of purified DNA was used for library preparation. The indexed libraries were hybridized and captured using the xGen^TM^ Custom Hyb Panel (IDT) targeting the 13 selected genes (STable 7) and sequenced on the DNBSEQ-G400RS (MGI Tech) in 150-bp paired-end mode. The consensus BAM file generation and mutation calling were performed as previously reported^53^. Briefly, sequence reads were aligned to the human reference genome (GRCh37) using BWA-MEM (v.0.7.17). Read families derived from the same parental molecule were identified by fgbio (v.1.4.0) and picard (v.2.24.0), based on the read start/end positions and the unique molecular identifiers (UMIs). Consensus reads were generated only for read families supported by ≥2 top reads and ≥1 bottom read; all other families were discarded. Somatic mutations were subsequently called using the Genomon pipeline (v.2.6.3). Modifications were applied to EBFilter (v.0.2.5) for accurate high-depth data processing and to GenomonMutationFilter (v.0.4.0). Called mutations were further filtered if they did not meet the following criteria: (i) consensus depth ≥3000×; (ii) VAFs ≥0.0012 and ≤0.20 in the sample; (iii) the variant read count in the paired normal control ≤1; (iv) strand ratio ≠0 or 1; (v) EBcall *P* value ≤10^-2.5^ using 20 duplex-sequencing data from non-paired peripheral blood or spleen; (vi) maximum variant reads detected in the same 20 non-paired control samples used for EBcall ≤1; (vii) the ratio of cruciform-derived reads among variant reads <0.5. Recurrent mutations observed in more than 20% of samples within a single case were further excluded.

### Validation of detected mutations

For WES validation, eight proximal tubule samples were newly collected from two donors using LCM. Lysates from each of these samples were divided into two aliquots, which were processed independently for DNA library preparation and sequencing. One BAM file was downsampled to match the average depth (49×) of the proximal tubule samples in this study, and mutation calling was performed. We then evaluated the resulting mutations at loci with sufficient sequencing depth (>75×) in the corresponding independent BAM files. A mutation was considered validated if it met all of the following criteria in the independent BAM file: (i) variant reads ≥2; (ii) VAF> 5 × VAF in the corresponding germline control samples; (iii) VAF > 2 × the maximum VAF observed across all other samples in the validation experiment (n=8). Consequently, the positive predictive value (PPV) for WES mutation calls was 94.8% (219/231 mutations).

For duplex sequencing validation, two independent experiments were performed using the same sample DNA, designated as the discovery and validation experiments (ED_Fig. 5e). Based on the mutation list obtained in the discovery experiment, a mutation was considered a true positive if it met all of the following criteria in the validation experiment: (i) variant reads ≥2; (ii) VAF > 5 × VAF in the corresponding germline control samples; (iii) VAF > 2 × the maximum VAF observed across all other samples in the validation experiment, excluding samples from the same punch (n=38). Consequently, the PPV for duplex sequencing mutation calls was 94.4% (117/124 mutations). The results of these validation experiments are summarized in STable 12.

### Analysis of significantly mutated genes

Significantly mutated genes were identified based on dN/dS using dndscv (v0.0.1.0)^55^. Shared mutations detected within the same tubule (in WES) or within the same punch (in duplex sequencing) were counted as single events. The dataset from duplex sequencing was analyzed by using a custom RefCDS object covering the targeted regions of the 13 genes. Genes with dN/dS>1 and *q*<0.1 were considered significantly mutated.

### Definition of highly reliable functional variants in RCC driver genes

To identify high-confidence functional mutations within genes significantly mutated in RCC, we applied specific filtering criteria to variants detected by WES and duplex sequencing. We defined “highly reliable functional variants” as mutations that were either truncating mutations in tumor suppressor genes, or variants registered as Tier 1, 2, or 3 in the COSMIC Cancer Mutation Census (v103)^56^.

### Copy number analysis

CNAs were evaluated on the basis of sequencing data using our in-house pipeline CNACS^4^. A total copy number (TCN) of >2.2 was defined as a gain and a TCN of <1.8 was defined as a loss. Copy number neutral loss of heterozygosity was identified based on a B-allele frequency <0.8 and a TCN between 1.8 and 2.2. All detected CNAs were manually curated. The results are summarized in STable 9. The fraction of cells harboring CNAs was estimated assuming a single-copy gain or loss; for autosomes, this was calculated as TCN - 2 for gains and 2 - TCN for losses. CNAs were classified as arm-level events if they affected ≥50% of a chromosome arm, while those spanning <50% were defined as focal events.

To estimate the proportion of cells harboring chromosome 18 loss (*C*minor), we performed allelic phasing using heterozygous single-nucleotide polymorphisms (SNPs). First, to identify heterozygous SNPs, mutation calling was performed on WES data from peripheral blood without matched controls. We applied the following filters to select reliable heterozygous SNPs on chromosome 18: (i) depth >15×; (ii) VAFs between 0.4 and 0.6; (iii) strand ratio ≠0 or 1; (iv) cruciform-derived reads <5. Finally, we randomly selected 80 heterozygous SNPs per patient. For each patient, we identified a representative sample with a clear allelic imbalance to phase the heterozygous SNPs on chromosome 18. Based on the VAFs in this reference sample, we assigned the A (retained) and B (lost) alleles and applied this phasing information to all other samples from the same patient. We calculated the mean VAF of the B allele (*VAF*minor) for each sample using the phased SNPs. *C*minor was then estimated as *C*minor = 2 × (0.5 - *VAF*minor).

### Calculation of mutant cell fraction and mutation clone size

Accurate estimation of the fraction of cells carrying a somatic mutation (mutant cell fraction; MCF) is confounded when the mutation resides in a chromosomal region affected by CNAs. For WES data analysis, to ensure the accuracy, we restricted the MCF analysis to mutations located in genomic regions with a neutral copy number state. Mutations within segments exhibiting copy number gains, losses, or uniparental disomies (UPDs) were excluded (587/33,791). For the retained mutations in neutral regions, MCFs were estimated as follows: MCF = 2 × VAF (for autosomes and the X chromosome in females); or MCF = VAF (for sex chromosomes in males). The mutation clone size was subsequently calculated as the product of the MCF and the corresponding LCM dissection area.

For mutations detected by duplex sequencing, a neutral copy number state was assumed for all targeted regions, and MCFs were calculated as 2 × VAF. Assuming a DNA content of 6.6 pg per diploid cell and a DNA recovery rate of 80%, the number of cells carrying the mutation was calculated using the following formula: (Number of cells) = (Total DNA (ng) / 0.8) × 2 × VAF / (6.6 × 10⁻³).

### Phylogenetic analysis

Phylogenetic trees were constructed for multiple samples obtained from single nephrons using somatic mutation and copy number data. Clonal hierarchies were inferred based on the cellular prevalence of each mutation cluster estimated using PyClone-VI (v0.1.1)^57^, applying the pigeonhole principle as previously described^5^. Briefly, if the sum of the cellular prevalences of two clusters exceeded 1.0, they were inferred to exhibit a nested relationship, with the cluster of lower prevalence defined as a subclone of the cluster with higher prevalence.

### Single-nucleus RNA-seq experiments

Single-nucleus suspensions were prepared following the protocol described by Muto et al.^58^, with minor modifications. Frozen kidney medulla specimens (1–2 mm³) were homogenized in 2 ml of lysis buffer (Nuclei EZ Lysis Buffer (Sigma-Aldrich) supplemented with 1 U/μl RNase Inhibitor (Roche)) using a Dounce Homogenizer (Kimble Chase). The suspension was transferred to a 15-ml tube, mixed with an additional 2 ml of lysis buffer, and incubated on ice for 5 min. The lysate was filtered through a 40-µm cell strainer (PluriSelect) and centrifuged at 500 × *g* for 5 min at 4 °C. The supernatant was discarded, and the pellet was resuspended in 4 ml of lysis buffer. After an additional 5-min incubation, the suspension was centrifuged again at 500 × *g* for 5 min at 4 °C and the supernatant was discarded. Finally, the nuclear pellet was resuspended in a nuclei suspension buffer (1× PBS containing 0.1% BSA and 0.1% RNase inhibitor) and filtered through a 10-µm cell strainer (PluriSelect). The single-nucleus suspension was assessed microscopically, and nuclei were counted using a Countess II Automated Cell Counter (Invitrogen). Single-nucleus RNA-seq libraries were prepared using the Chromium GEM-X Single Cell 3’ Kit v4 (10x Genomics) targeting 20,000 nuclei per sample, following the manufacturer’s instructions. Sequencing was performed on the DNBSEQ-G400RS platform (MGI Tech) using a custom protocol (Read 1: 28 cycles; Read 2: 90 cycles).

### Single-nucleus RNA-seq analysis

Barcode processing and gene expression quantification were performed using 10x Genomics Cell Ranger (v9.0.1)^59^ with default parameters and the GRCh38 reference genome. To correct for ambient RNA contamination, raw count matrices were processed using SoupX (v1.6.2)^60^. Subsequent quality control and downstream analyses were conducted using Seurat (v5.1.0)^61^. Low-quality nuclei were filtered out based on the following criteria: mitochondrial RNA content >5%, and nFeature_RNA or nCount_RNA values falling below sample-specific thresholds (STable 13). Doublets were identified and removed using DoubletFinder (v2.0.4)^62^. Furthermore, nuclei exhibiting anomalously high nFeature_RNA and nCount_RNA values were excluded to eliminate potential homotypic doublets (STable 13). The filtered Seurat objects were normalized and scaled using the SCTransform function. Datasets from all cases were integrated using Harmony (v1.2.3)^63^ within the Seurat framework. Unsupervised clustering was performed at a resolution of 0.5 based on the first 20 principal components. UMAP embedding was used for data visualization. Cell type annotation was performed manually based on the expression of established marker genes^45^ (ED_Fig. 8c).

### Copy number analysis of single-nucleus RNA-seq data

To infer CNAs from single-nucleus gene expression profiles, we used inferCNV (v1.2.0)^33^. The collecting duct cell population, comprising principal cells (CD-PCs) and intercalated cells (CD-ICs), was subsetted from the integrated dataset. The inferCNV pipeline was run independently for each patient, using raw count matrices and cell type annotations as inputs with the following parameters: cutoff = 0.1, cluster_by_groups = TRUE, denoise = TRUE, and leiden_resolution = 0.01. We manually annotated the CNA status for each subcluster identified in the resulting dendrogram. CNAs consistently observed across all clusters of CD-PCs or CD-ICs were regarded as reflections of lineage-specific expression profiles and were excluded from the CNA annotations. Subclusters accounting for less than 2.5% of the total cells within each population (CD-PC or CD-IC) were labeled as ambiguous due to low confidence in CNA determination. After assigning CNA status to the metadata, the collecting duct cells from all cases were re-integrated using Harmony and re-clustered with Seurat. Clusters 3 and 8 were enriched for cells exhibiting a gain of chromosome 13q (SFig. 5a-b). Notably, these clusters showed concurrent high expression of markers specific to both the distal straight tubule and the collecting duct (SFig. 5b), suggesting doublets or ambient RNA contamination that had not been removed during initial filtering. Consequently, Clusters 3 and 8 were excluded, and the inferCNV pipeline was repeated on the filtered dataset, with the final output shown in ED_Fig. 9a.

### Pseudobulk differential expression analysis and gene set enrichment analysis

To investigate transcriptional changes associated with CNAs in CD-PCs, we performed a pseudobulk differential expression analysis. Pseudobulk expression profiles were generated by aggregating raw gene counts for CD-PCs from each patient, stratified by their CNA status using the AggregateExpression function in Seurat. Differential expression analysis was performed using DESeq2 (v1.50.2)^64^, comparing nuclei with specific CNAs against wild-type nuclei. Specifically, we analyzed chromosome 18 loss using data from 4 patients and chromosome 9 loss using data from 2 patients. To account for inter-patient variability, patient identity was included in the design formula as follows: design = ∼ patient + CNA_status. Gene set enrichment analysis was performed using fgsea (v1.36.0)^65^. All genes were pre-ranked based on the Wald statistic derived from the DESeq2 analysis, and enrichment was tested against the MSigDB Hallmark gene sets^66^.

### Quantification of the stromal proportion

The proportion of stroma was quantified using HALO software (Indica Labs). For each H&E-stained slide, parenchymal regions (comprising glomeruli, renal tubules, and large vessels) and stromal regions (defined as areas exclusive of parenchyma) were annotated, with a median of 606 training regions outlined across both areas per case. The MiniNet module was subsequently trained for each individual case with the following configurations: resolution = 0.25 μm/px and minimum object size = 100 μm^2^. Using the trained classifier, the classification was expanded to the entire sampling area (median analysis area = 4.8 mm^2^). The proportion of stroma was calculated as the ratio of the annotated stromal area to the total analyzed area.

## Statistical analysis

Statistical analyses were performed using R (v.4.5.0). All *P* values were two-sided. For group comparisons, the Wilcoxon signed-rank test was used. To compare differences in proportions between groups, Fisher’s exact test was performed, and the odds ratio with 95% confidence interval (CI) was calculated. Multiple testing was corrected using the Benjamini-Hochberg method.

The associations between donor age and the number of somatic mutations detected by NanoSeq were evaluated using linear regression models without assuming a zero intercept. For individuals with multiple samples, the mean values were used as a single representative value. To compare the slopes of the regression lines between the two groups, we tested the significance of the interaction term in a linear regression model.

The association between donor age and the number of CNAs per sample was evaluated using a generalized linear mixed-effects model (GLMM) with a Poisson distribution and a log link function. Donor identity was included as a random effect.

For the comparison of median mutation clone sizes in proximal tubules (PCT and PST) between young (26 years old) and aged (76 years old) donors, 95% CIs were estimated using bootstrap resampling of samples with 10,000 iterations. To identify factors associated with mutation size, multivariate analysis was conducted using a GLMM assuming a Gamma distribution with an identity link. We incorporated donor identity as a random effect to model median mutation size per 1,000 μm^2^ against the presence of CNAs and highly reliable functional variants in RCC driver genes, and age in 10-year increments across nephron segments.

For the estimation of driver mutation and CNA frequencies, as well as the cell fraction of CNAs from WES analysis, we calculated adjusted values using a resampling approach to prevent overrepresentation of samples derived from the same tubule. The total number of unique tubules was defined as N = a + b, where a is the number of solitary tubules and b is the number of tubule groups from which multiple contiguous samples were collected. For each of the b groups, one sample was randomly selected and combined with solitary samples to generate a representative dataset of N samples. The frequency of driver mutations or CNAs, as well as the cell fraction for each CNA were then calculated within this dataset. This resampling and calculation process was repeated 1,000 times. The final adjusted frequency and mean cell fraction were reported as the mean of these 1,000 replicates, with error bars representing the 95% CI determined by the percentile method.

The associations between the chromosome 18-deficient cell fraction and the proportion of stroma were evaluated using linear regression models without assuming a zero intercept. For individuals subjected to contiguous sampling, the adjusted cell fractions were used.

## Data and Availability

All raw sequencing data generated in this study will be deposited in the European Genome-phenome Archive (EGA) under study accession EGAS50000002066, with dataset accession numbers EGAD50000003153 (NanoSeq); EGAD50000003035, EGAD50000003036, and EGAD50000003037 (WES); EGAD50000003087 (WGS); EGAD50000003022 and EGAD50000003023 (DuplexSeq); and EGAD50000002967 (snRNA-seq). All other data are available from the corresponding author upon reasonable request.

## Code Availability

The code for snRNA-seq analysis, the custom RefCDS file used for dN/dS analysis of duplex sequencing, and the SNP/noise masks for NanoSeq analysis are available via Zenodo (DOI: 10.5281/zenodo.18726765).

## Acknowledgements

We express our gratitude to the donors and their families for their donation of biological specimens, and we thank the clinicians who provided these biospecimens. We express our sincere gratitude to Dr. Benjamin D. Humphreys for his insightful comments and valuable discussion regarding this work. We are grateful to the Single-cell Genome Information Analysis Core (SignAC) at WPI-ASHBi, Kyoto University, and the Division of Histological Study, Center for Anatomical Studies, Graduate School of Medicine, Kyoto University for technical support. We also thank T. Sakurai, Y. Indo, K. Onishi, T. Shirahari, A. Ryu, N. Hakoda, E. Gunshima, and M. Ushijima for their technical assistance, and Y. Tobimatsu for valuable suggestions on the manuscript. Fig. 1a was created with BioRender.com. This work was supported by the Japan Agency for Medical Research and Development (AMED): The Moonshot Research and Development Program (grant nos. JP22zf0127009 to S.O. and JPMJMS2022-25 to N.K.) and the Project for Promotion of Cancer Research and Therapeutic Evolution (JP24ama221530); the Japan Society for Promotion of Science (JSPS): Specially Promoted Research (grant no. JP24H00009) and KAKENHI (JP24K23455 to K.I.); the Japan Science and Technology Agency (JST): Fusion Oriented Research for Disruptive Science and Technology (FOREST) Program (grant no. JPMJFR215V to N.K.); Takeda Science Foundation (to S.O. and N.K.); Princess Takamatsu Cancer Research Fund (to N.K.); The Naito Foundation (to N.K.); Inoue Foundation for Science (to N.K.); and Nanpuh Hospital (to S.O.).

## Author Contributions

K.I., N.K., M.E. and S.O. designed the study. G.F., D.T., S.N., T.T., S.U., T.S., S.Kobayashi, J.M., S.G., T.M., K.M., E.K., J.I., Y.Oda, R.N., T.R.K., M.T., M.H., M.S. and M.E. provided patient samples and collected clinical data. G.F., D.T. and Y.Oda performed histological diagnosis. K.I., T.Nakagawa, G.F., D.T., S.N., T.T., S.U., T.S., S.Kobayashi, J.M., S.G., T.M., K.M., E.K., J.I., R.N., T.R.K., M.T., M.H. and M.S. performed sample preparation. K.I., N.K., T.H., K.W., S.Kawaguchi, H.M., Y.I., T.Nishimura, R.S. and T.Nakagawa developed methodology. K.I., N.K., T.H., K.W., S.Kawaguchi, H.M., Y.I., T.Nishimura, T.Noda, M.M., M.M.N., R.S., Y.Ochi and S.M. performed bioinformatics. K.I., N.K., M.M.N. and S.O. wrote the manuscript. M.E. and S.O. jointly lead the study. All authors reviewed and approved the manuscript.

---

**Extended Data Figure 1.**
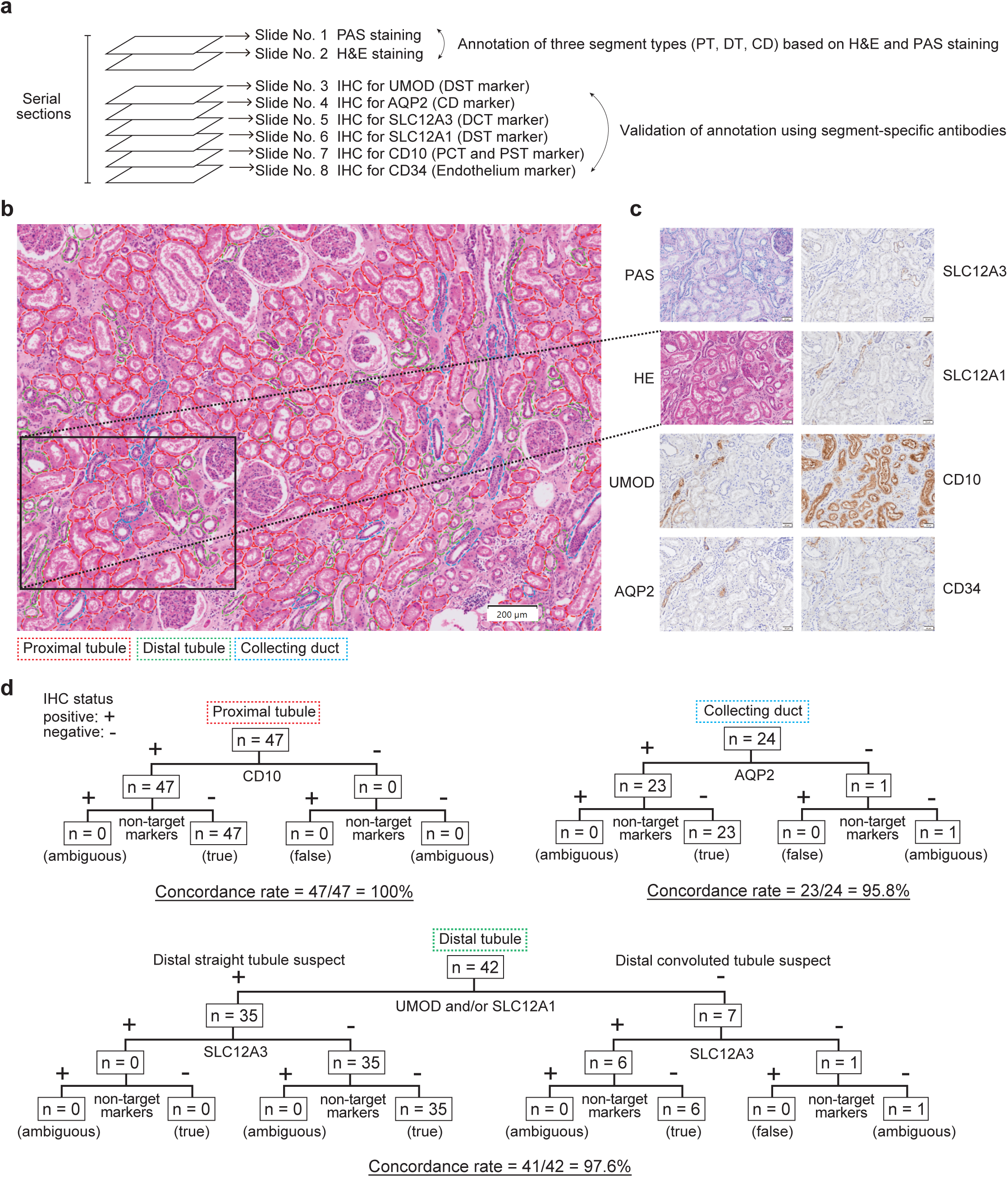
Validation of nephron segment annotation by immunohistochemistry (IHC). a, Experimental design. Serial sections were prepared for H&E, PAS and IHC staining. Three segment types (PT: proximal tubule, DT: distal tubule, and CD: collecting duct) were annotated based on H&E and PAS staining, followed by validation using segment-specific antibodies. **b,** Annotation of the three segments overlaid on the H&E image. **c,** IHC staining using segment-specific antibodies on serial sections. **d,** Concordance between H&E/PAS-based annotation and validation by IHC using segment-specific antibodies.

**Extended Data Figure 2.**
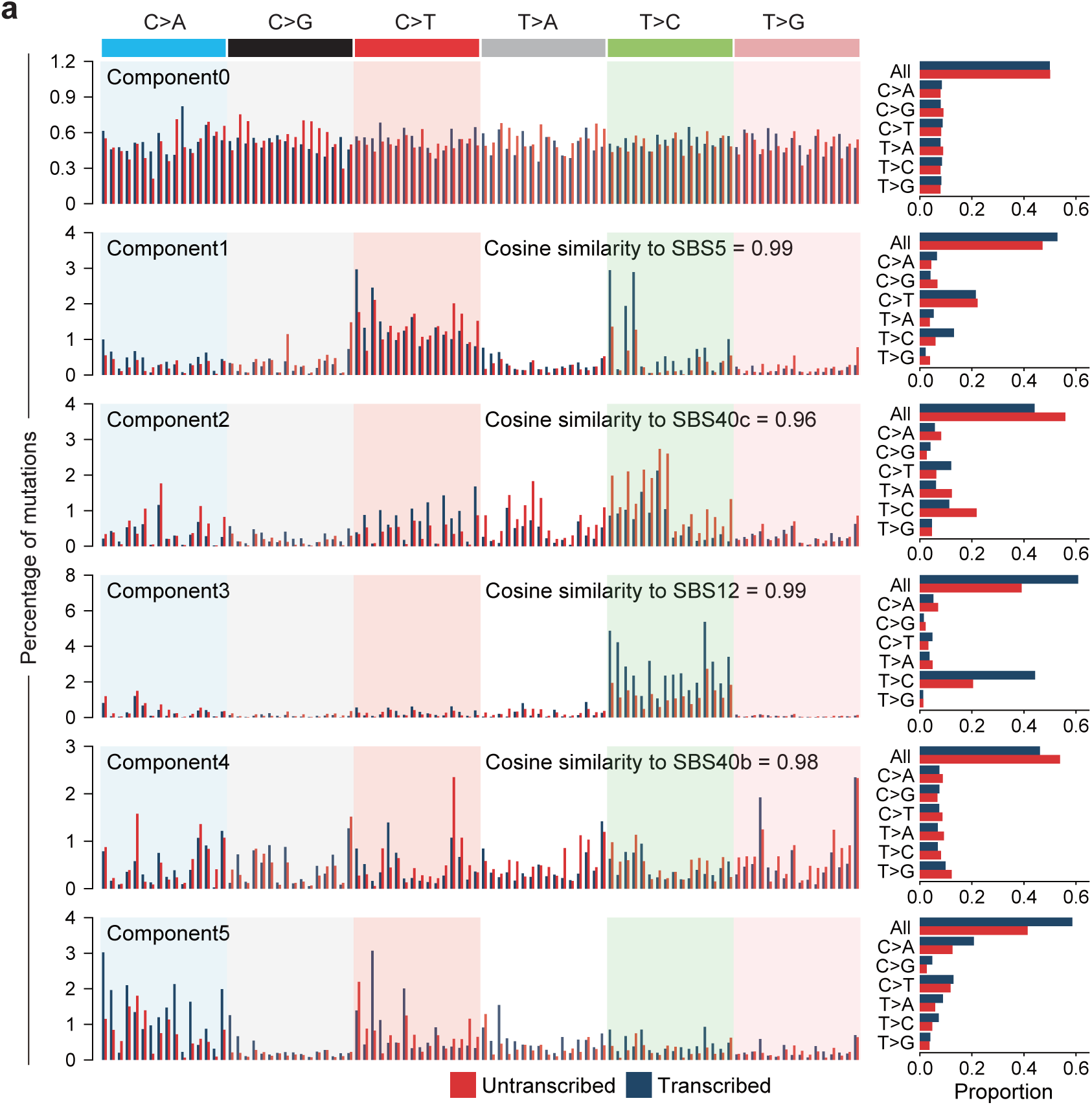
Transcriptional strand asymmetry of SBS signatures in normal nephrons. a, Five *de novo* substitution signatures showing transcriptional strand bias were extracted using HDP. Cosine similarities to signatures in Fig. 2a were calculated after collapsing the 192 substitution classification into the standard 96 substitution classification. Only cosine similarities ≥0.9 are shown.

**Extended Data Figure 3.**
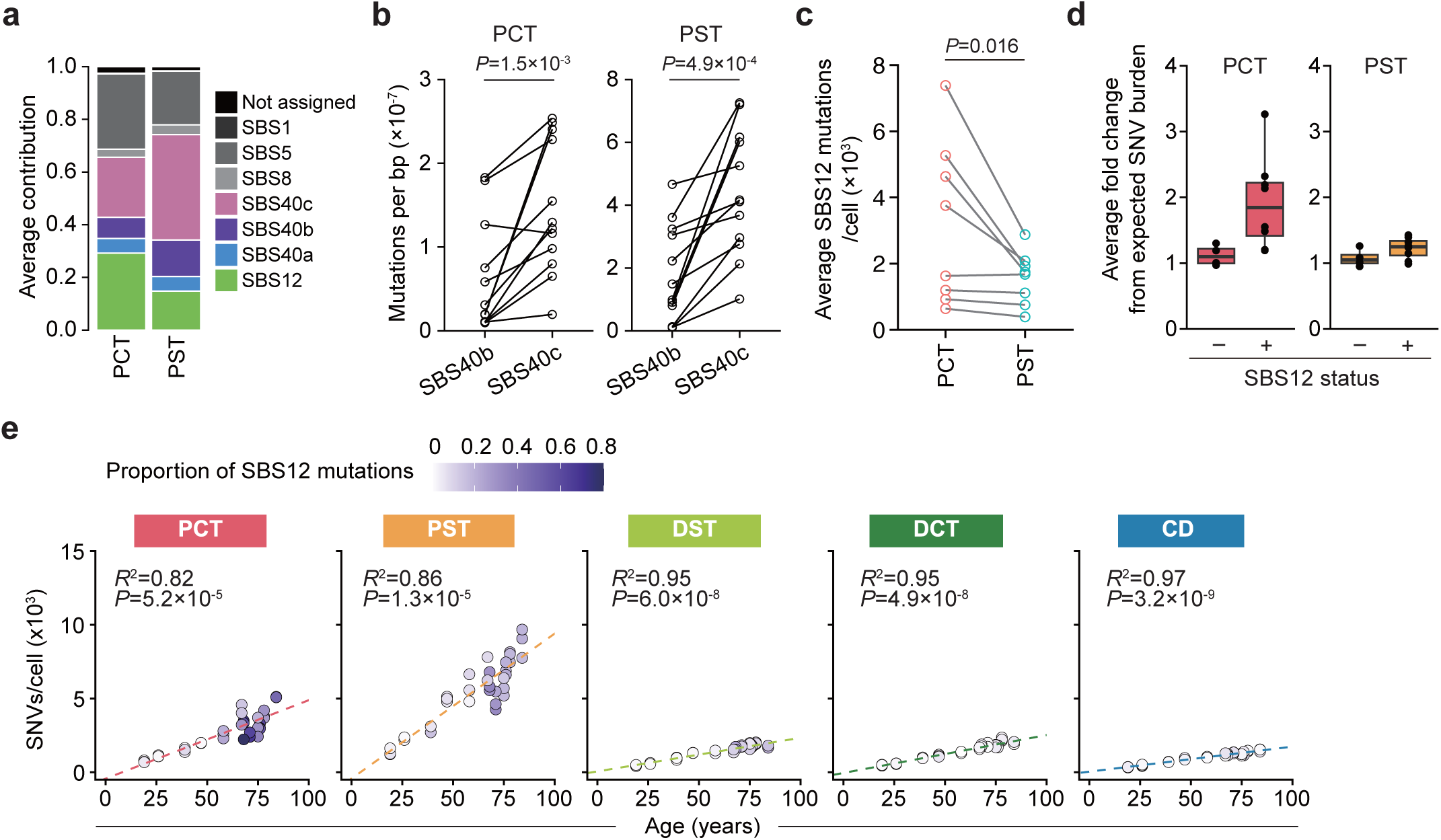
Mutational signatures in PCT and PST. a, Average contribution of SBS signatures in PCT and PST. **b,** Paired comparison of mean burdens of SBS40b and SBS40c mutations per base pair between PCT and PST. *P* value from the Wilcoxon signed-rank test. **c**, Paired comparison of mean SBS12 mutations per cell between PCT and PST in SBS12-positive cases (n=8). *P* value from the Wilcoxon signed-rank test. **d,** Average fold change of observed versus age-expected SNV burden in samples with and without SBS12 exposure. **e,** Correlation between mean numbers of SNVs/cell (excluding SBS12 contribution) and donor age in indicated nephron segments. Content of SBS12 mutations is shown as a color gradient. *R*^2^ and *P* values were obtained using a linear regression model.

**Extended Data Figure 4.**
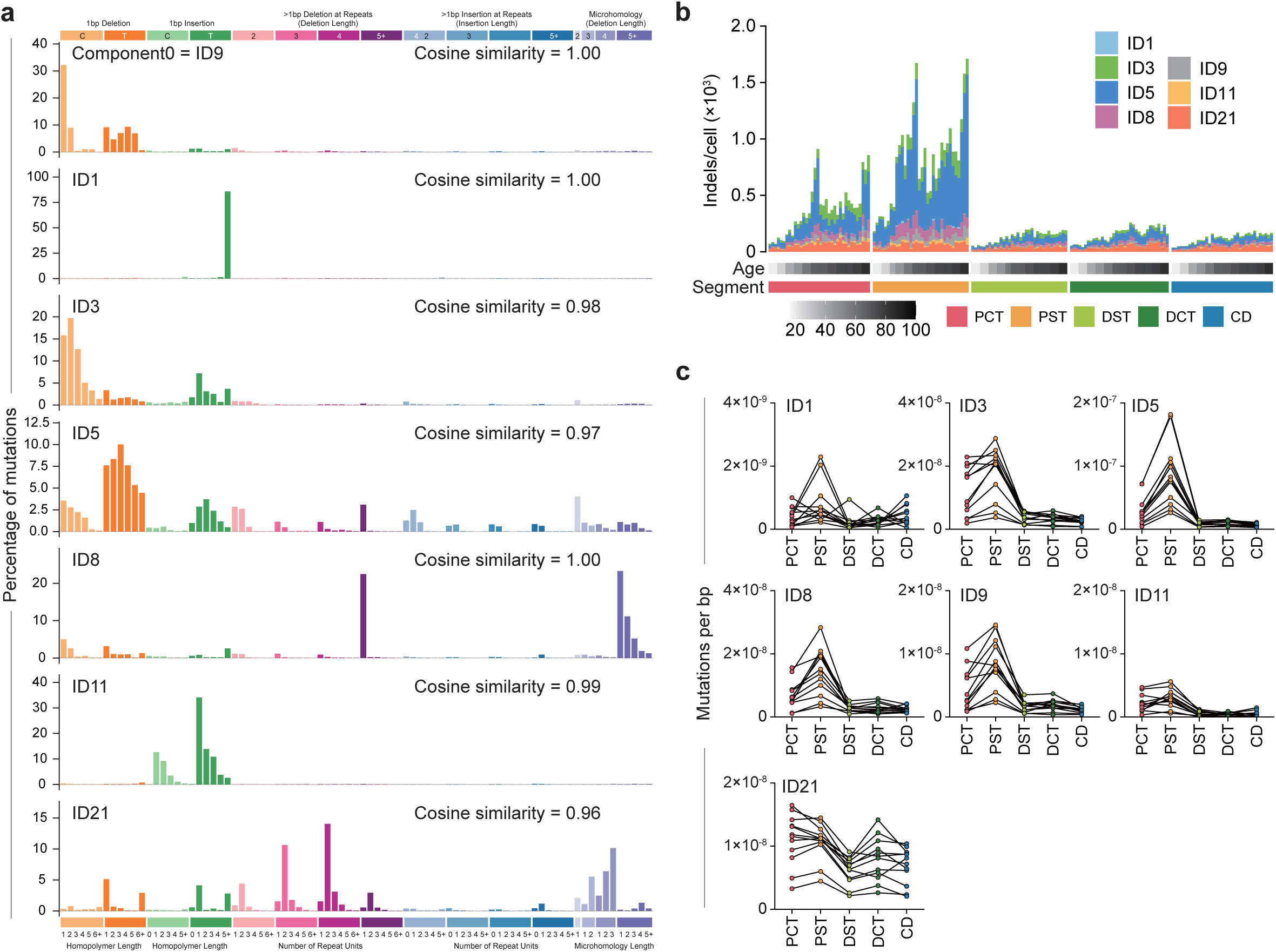
ID signatures in normal nephrons. a, Barplots show 83 mutational patterns for ID signatures extracted using HDP. Cosine similarities were calculated against the COSMIC ID signature catalogue. **b,** Landscape of ID signatures across segments. **c,** Average mutation burden of ID signatures across segments and donors. Lines connect samples from the same donor.

**Extended Data Figure 5.**
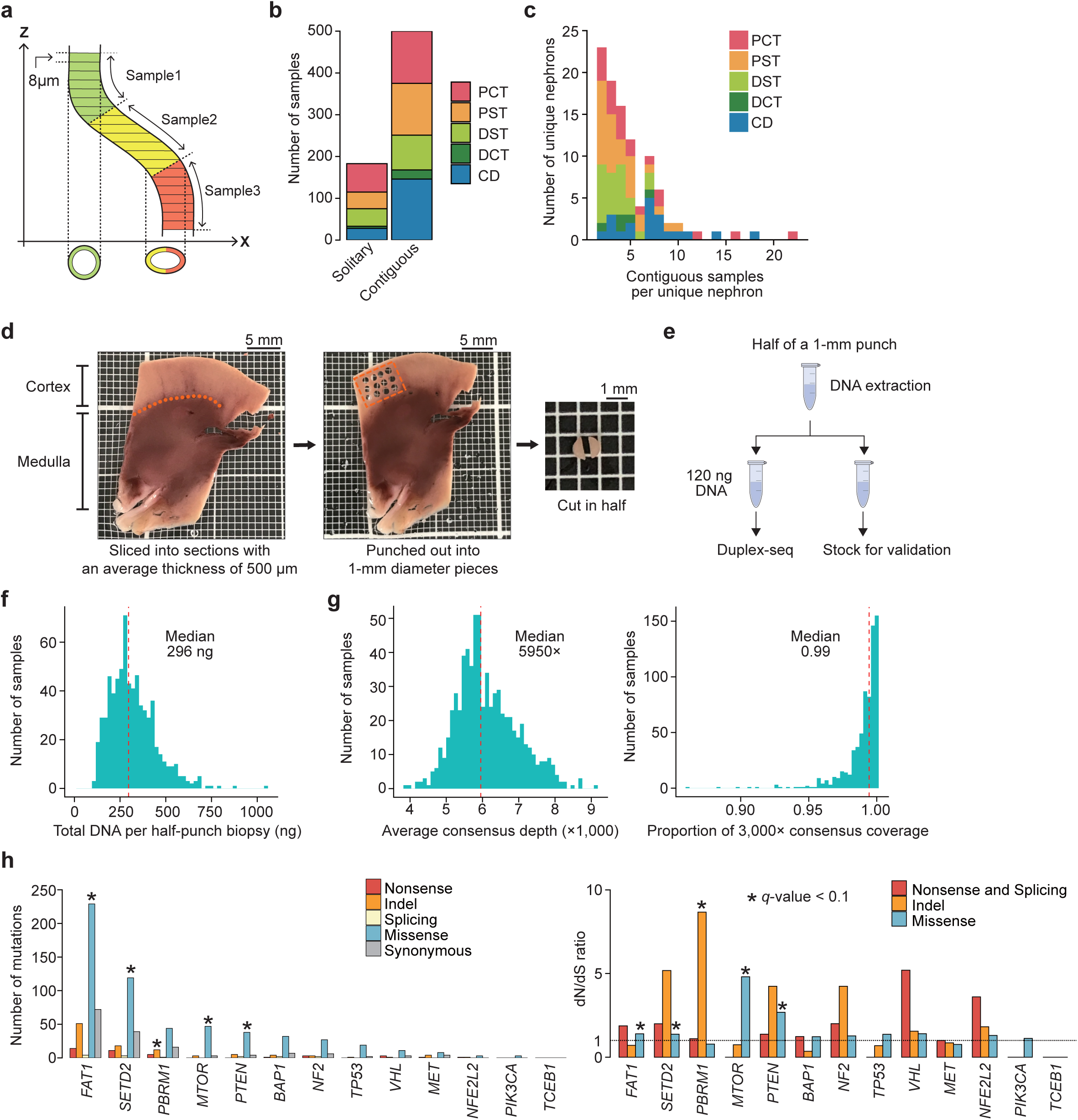
Sampling strategy for WES and duplex sequencing, along with quality metrics and mutational profile. a, Schematic of contiguous sampling of a single nephron from serial sections for WES. **b,c,** Number of solitary and contiguous WES samples (b) and a histogram of contiguous samples per unique nephron (c). **d,e**, Sample preparation workflow for duplex sequencing, in which half-punch biopsies (1 mm in diameter; mean volume 0.2 mm³) were collected from the cortex and medulla of kidney slices (d) and 120ng of extracted DNA was subjected to duplex sequencing, with the remainder reserved for validation (e). **f**, Total DNA per half-punch biopsy. **g,** Average consensus depth (left) and the proportion of the target region covered at ≥3,000× consensus depth (right) for duplex sequencing. **h,** Number of mutations detected by duplex sequencing (left) and their dN/dS ratios (right). Asterisks indicate mutation types with *q*<0.1 in dN/dS analysis.

**Extended Data Figure 6.**
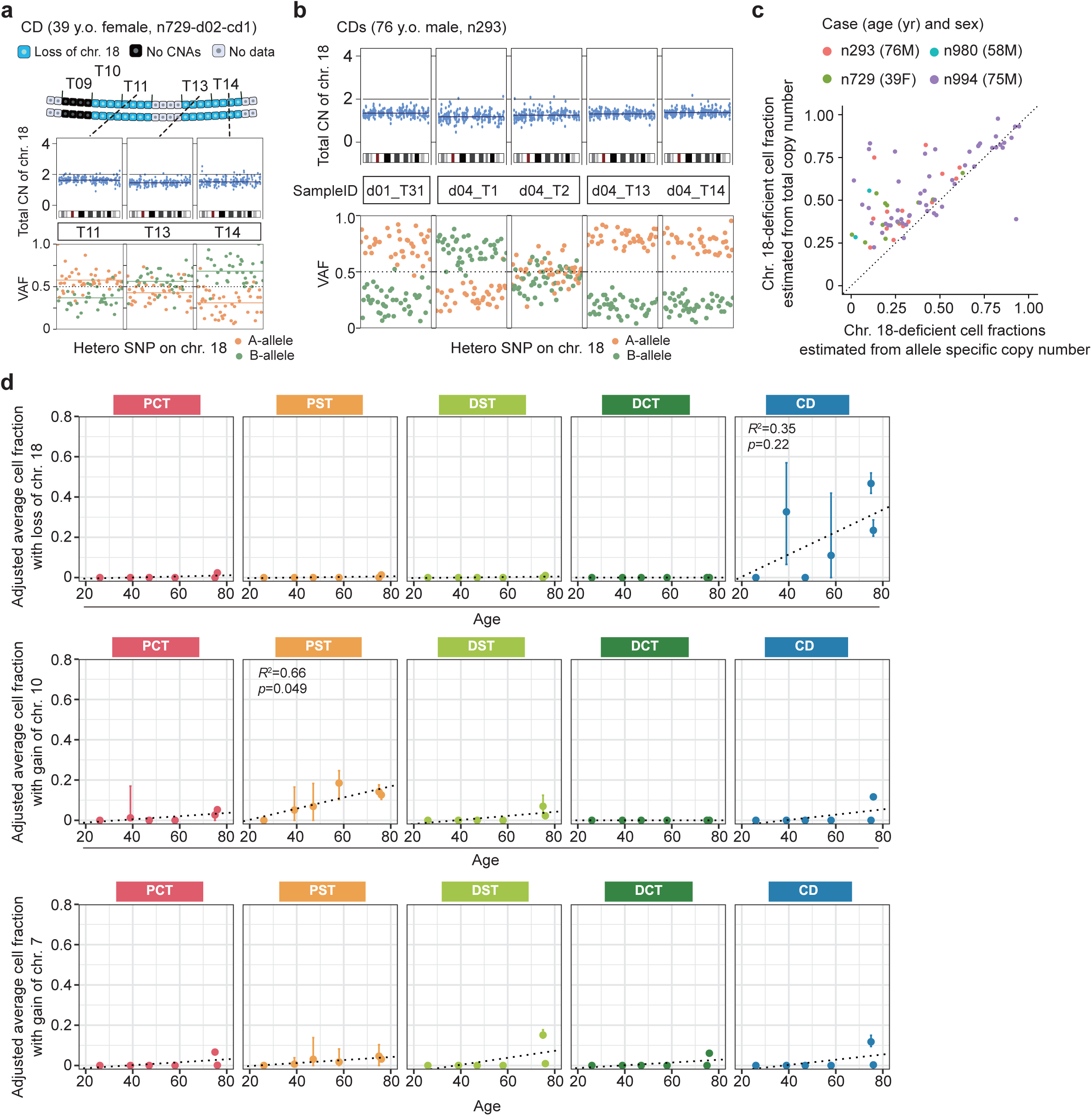
Copy number alterations in normal nephrons. a, Two distinct clones carrying chromosome 18 loss found along a CD segment in a 39-year-old female donor (top). Bottom panels show total-copy number profile and A- and B-allele frequencies at heterozygous SNPs on chromosome 18 in three samples. A horizontal bar is included in the bottom panels to indicate the mean VAF for the A-allele and B-allele, respectively. **b,** Total copy numbers and A- and B-allele VAFs at heterozygous SNPs on chromosome 18 for samples with a total copy number ≤ 1.5 from a 76-year-old male donor. Samples from the same nephron are enclosed in boxes. **c,** Chromosome 18-deficient cell fractions estimated from total copy numbers (y-axis) and allele-specific copy numbers (x-axis) **(Methods)**. **d,** Average cell fraction of recurrent CNAs in each nephron segment, adjusted for contiguous sampling **(Methods).**

**Extended Data Figure 7.**
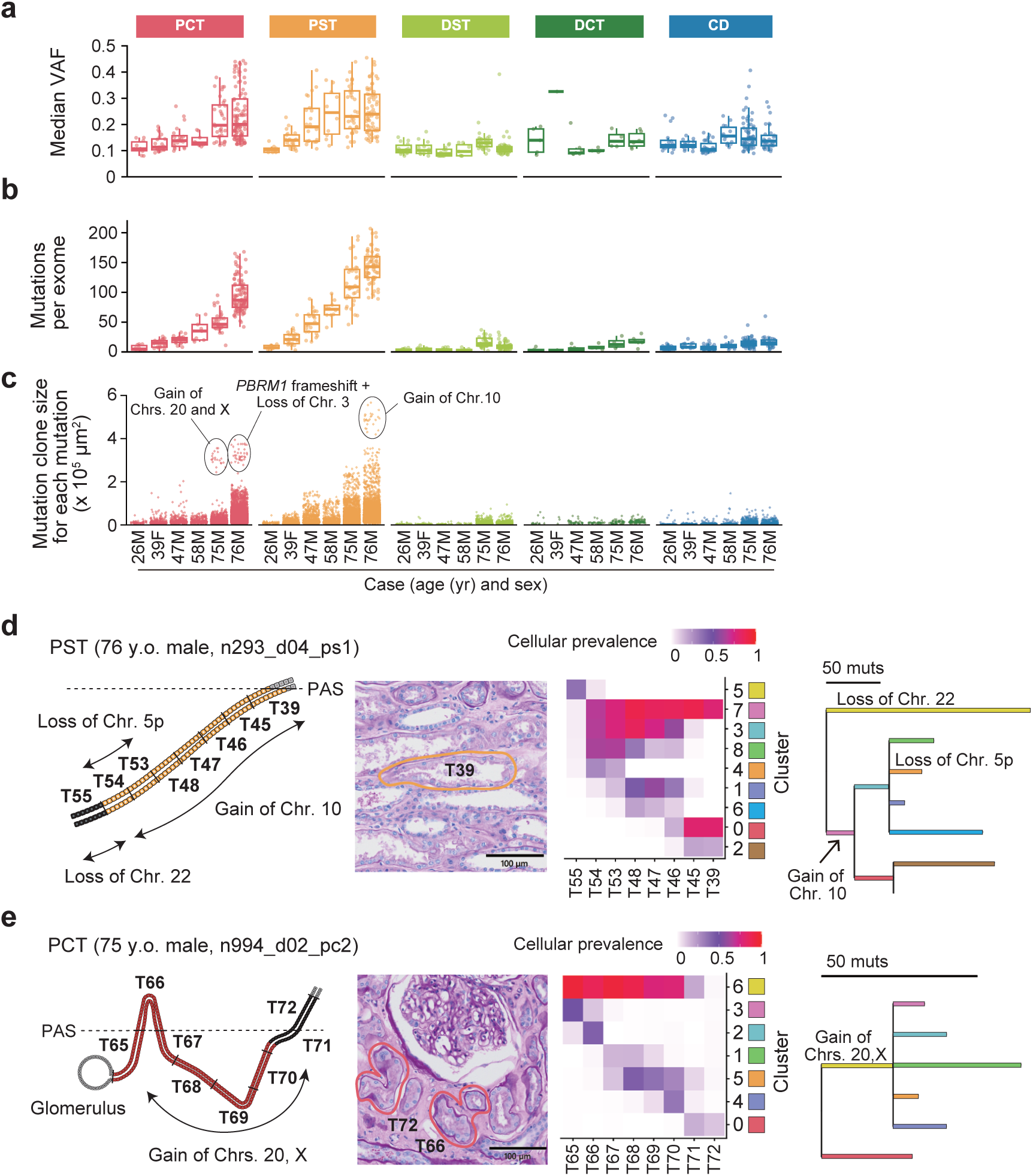
Mutation clone size and extensive clonal expansion in aged nephrons from contiguous LCM samples. a,b, Median VAFs (**a**) and numbers (**b**) of SNVs and indels detected by WES across 683 LCM samples from five nephron segments are plotted for each sample and donor. Box plots indicate the median (centre line), interquartile range (box) and 1.5 × IQR (whiskers). **c,** Mutation clone size per mutation; sizes are aggregated for any mutations spanning multiple samples. **d,e,** Representative examples of extensive clonal expansion in aged nephrons within the PST (**d**) and PCT (**e**). Panels display schematic nephron architecture, PAS staining at the indicated level, cellular prevalence of mutational clusters inferred by PyClone-VI, and corresponding phylogenetic trees.

**Extended Data Figure 8.**
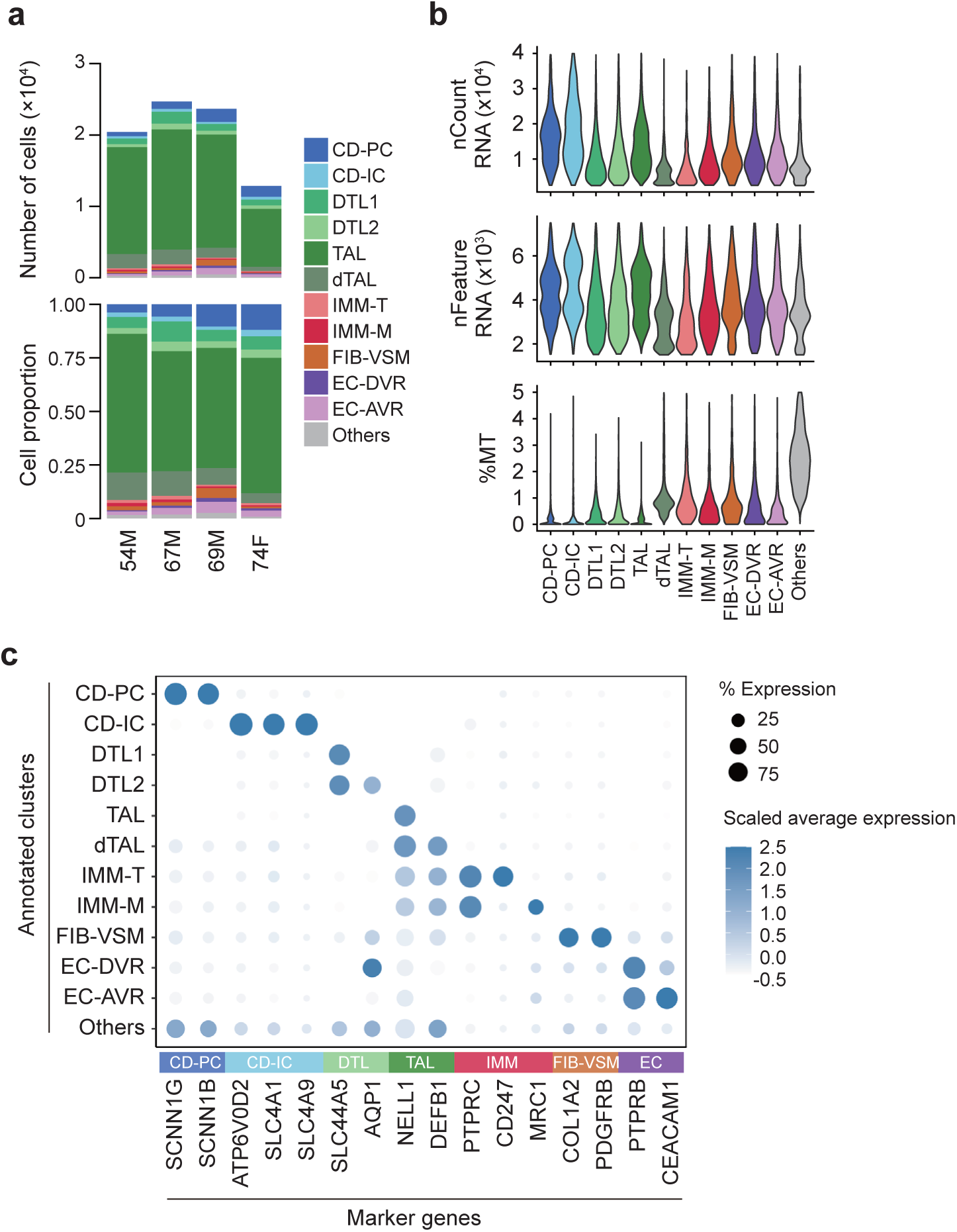
Quality metrics and cell annotation for snRNA-seq. a, Cell counts and proportions for each annotated cluster per patient. CD-PC, principal cell of collecting duct; CD-IC, intercalated cell of collecting duct; DTL1, descending thin limb cell type1; DTL2, descending thin limb cell type2; TAL, thick ascending limb cell; dTAL, degenerative thick ascending limb cell; IMM-T, T cell; IMM-M, macrophage; FIB-VSM, fibroblast and vascular smooth muscle cells; EC-DVR, descending vasa recta endothelial cell; EC-AVR, ascending vasa recta endothelial cell. **b,** Quality metrics summary of each annotated cluster. **c,** Expression of marker genes for each annotated cluster.

**Extended Data Figure 9.**
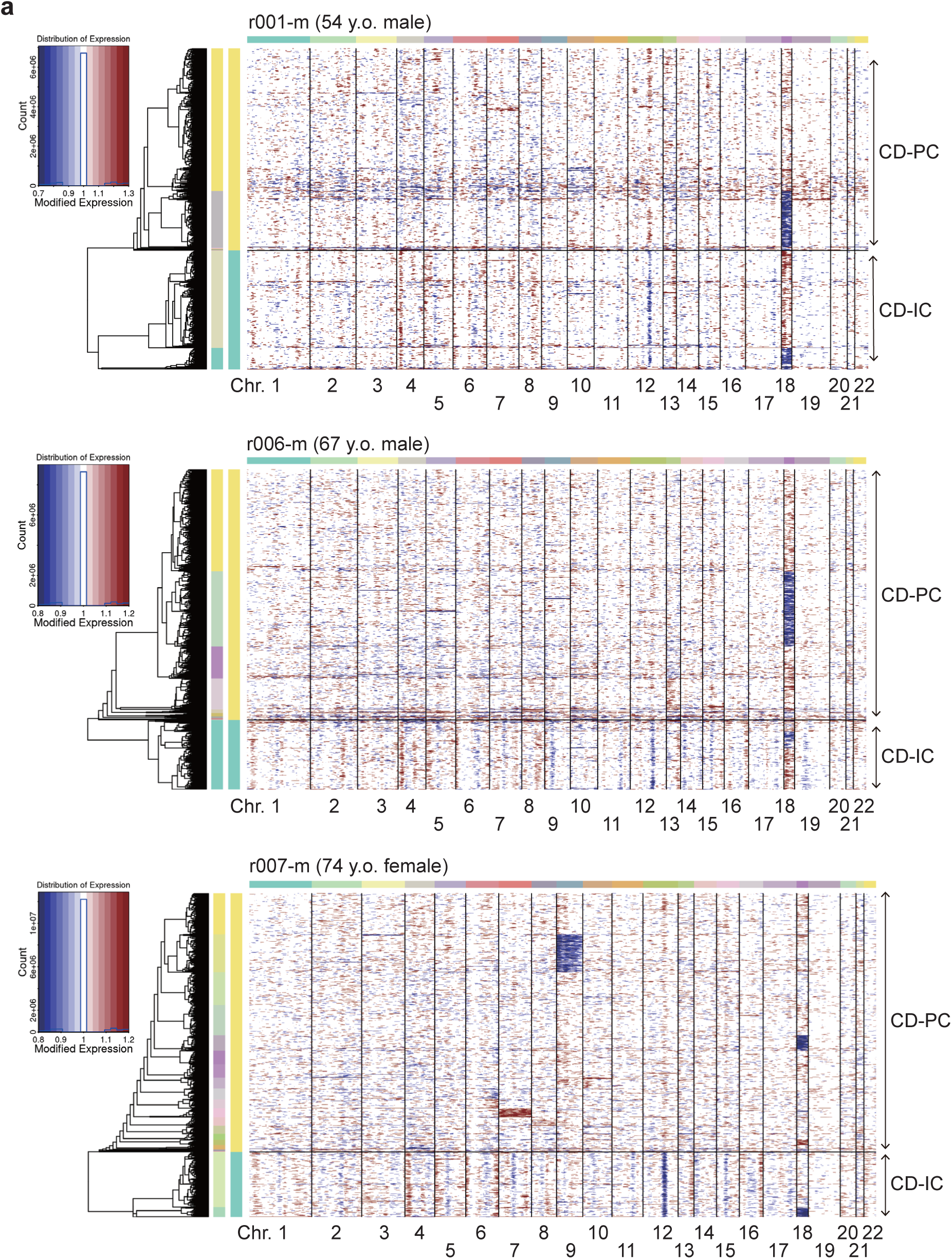
InferCNV analysis of snRNA-seq data. a, InferCNV heatmaps showing CNAs in CD-PCs and CD-ICs for medullary samples from the three donors not shown in Figure 5b: r001-m (54-year-old male), r006-m (67-year-old male), and r007-m (74-year-old female). Red indicates chromosomal gains, and blue indicates losses.

**Extended Data Figure 10.**
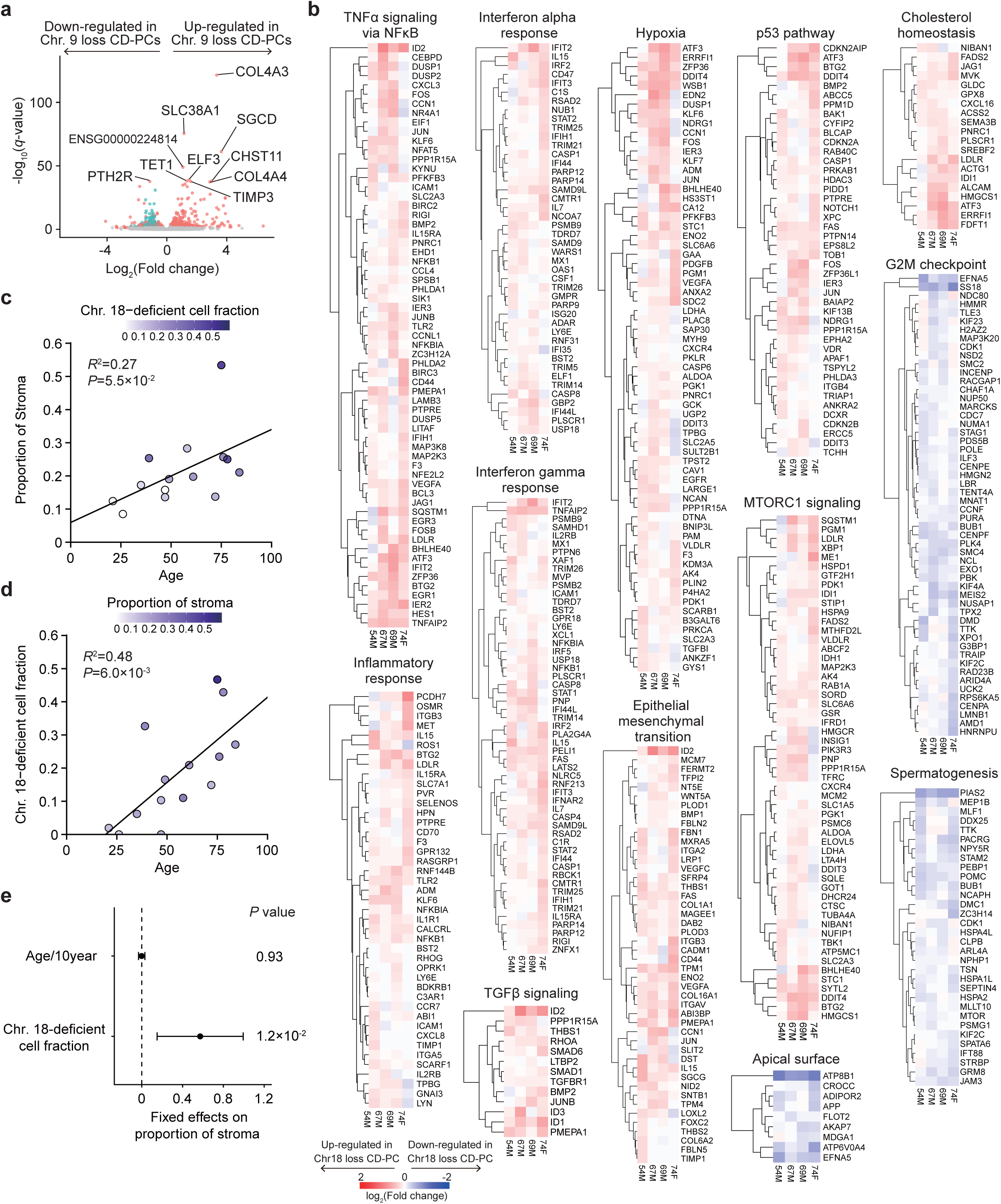
Impact of chromosomal losses on gene expression and stromal proportions in CD-PCs. a, Volcano plot of DEGs identified in pseudo-bulk analysis comparing CD-PCs with and without chromosome 9 loss. Significant DEGs (*q*<0.05 and |log2FoldChange|>0.25) are colored red or green, with genes on chromosome 9 highlighted in green. **b,** Heatmap of patient-specific log2 fold changes of leading-edge genes in MSigDB Hallmark gene sets significantly enriched in chromosome 18-loss CD-PCs compared to wild-type (WT). **c**,**d**, The proportion of stroma (c) and the chromosome 18-deficient cell fraction (d) are plotted against donor age, with fitted regression lines. *R*^2^ and *P* values were obtained using a linear regression model. For patients with contiguous samples, adjusted cell fractions were used. **e**, Estimated fixed effects on proportion of stroma and *P* values calculated using a multiple linear regression model. Explanatory variables included age (per 10 years) and chromosome 18-deficient cell fraction. Error bars indicate 95% confidence intervals.

**Supplementary Figure 1.**
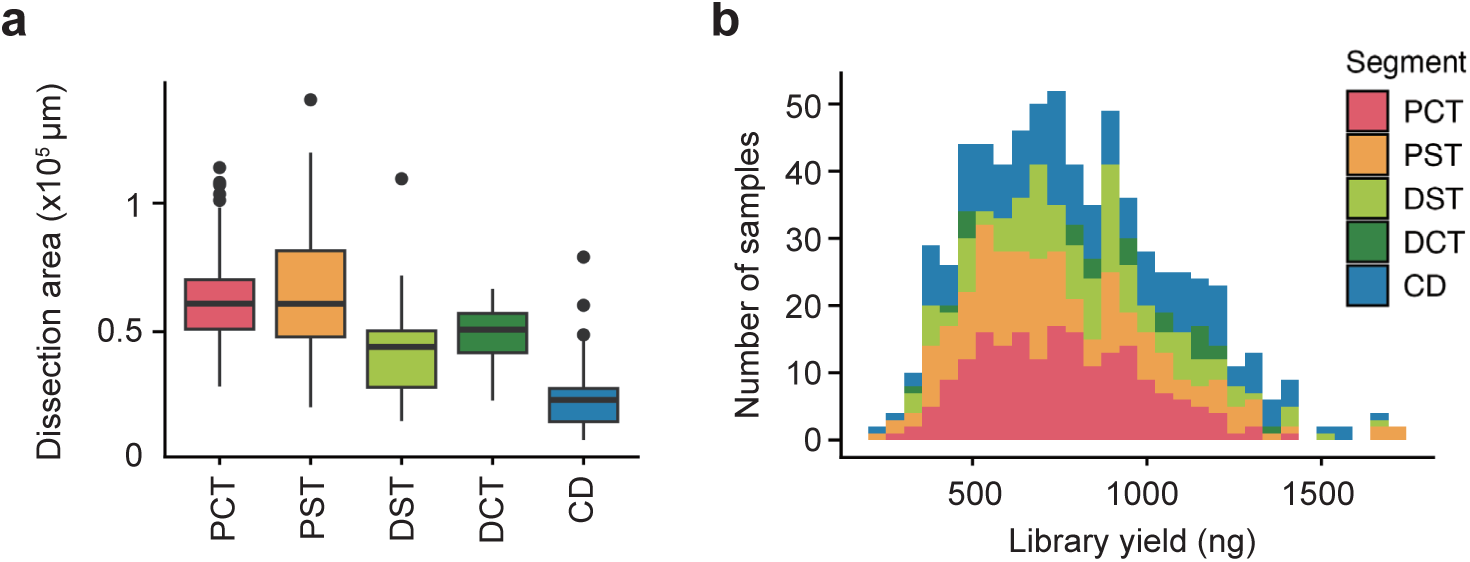
Quantitative input control for LCM samples. a,b, Distribution of LCM dissection area (a) and library yield (b) for each nephron segment across 683 WES samples.

**Supplementary Figure 2.**
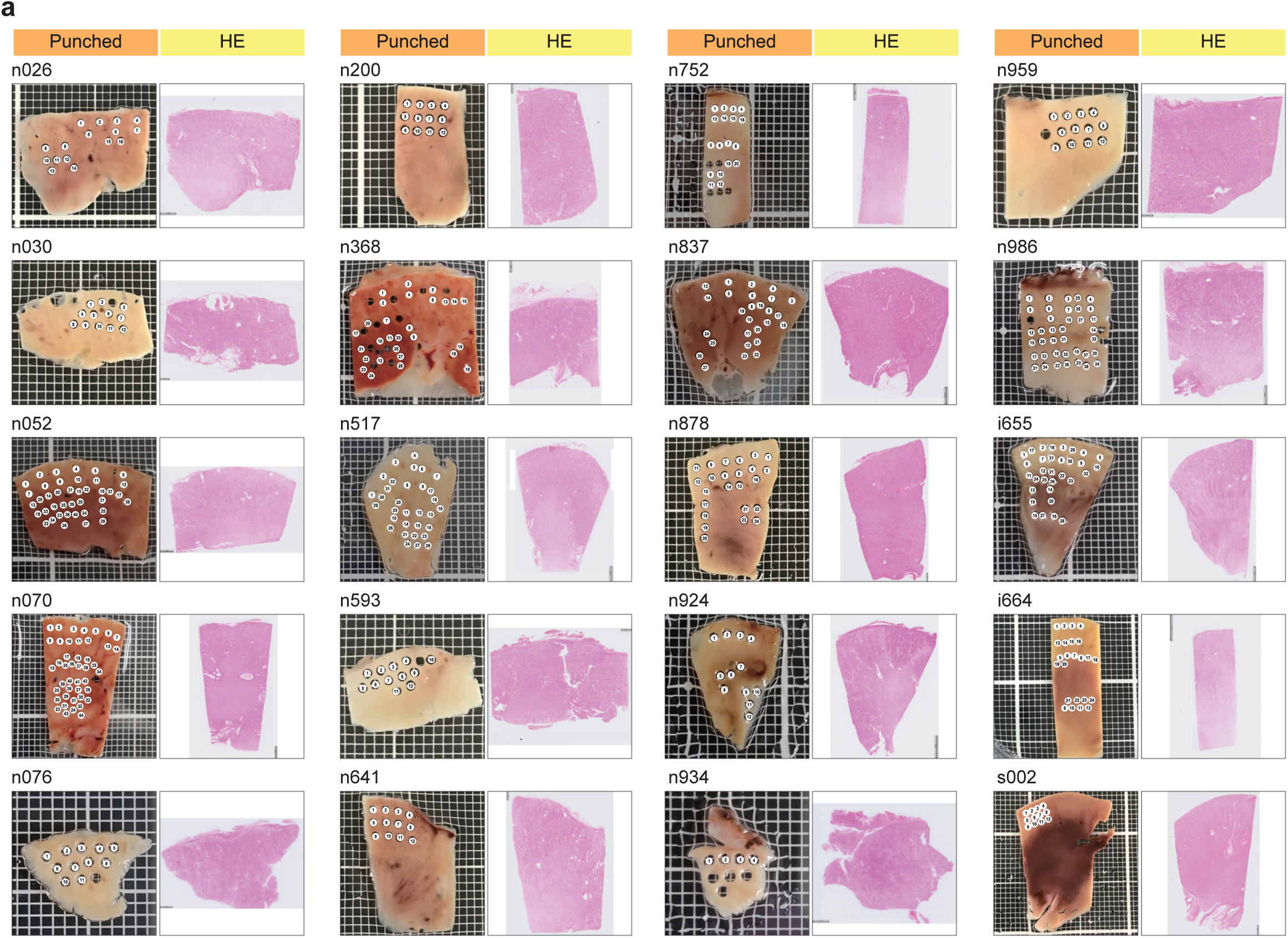
Overview of kidney samples and histology for duplex sequencing. a, Images of bulk samples and adjacent H&E-stained sections for the 20 donors analyzed by duplex sequencing.

**Supplementary Figure 3.**
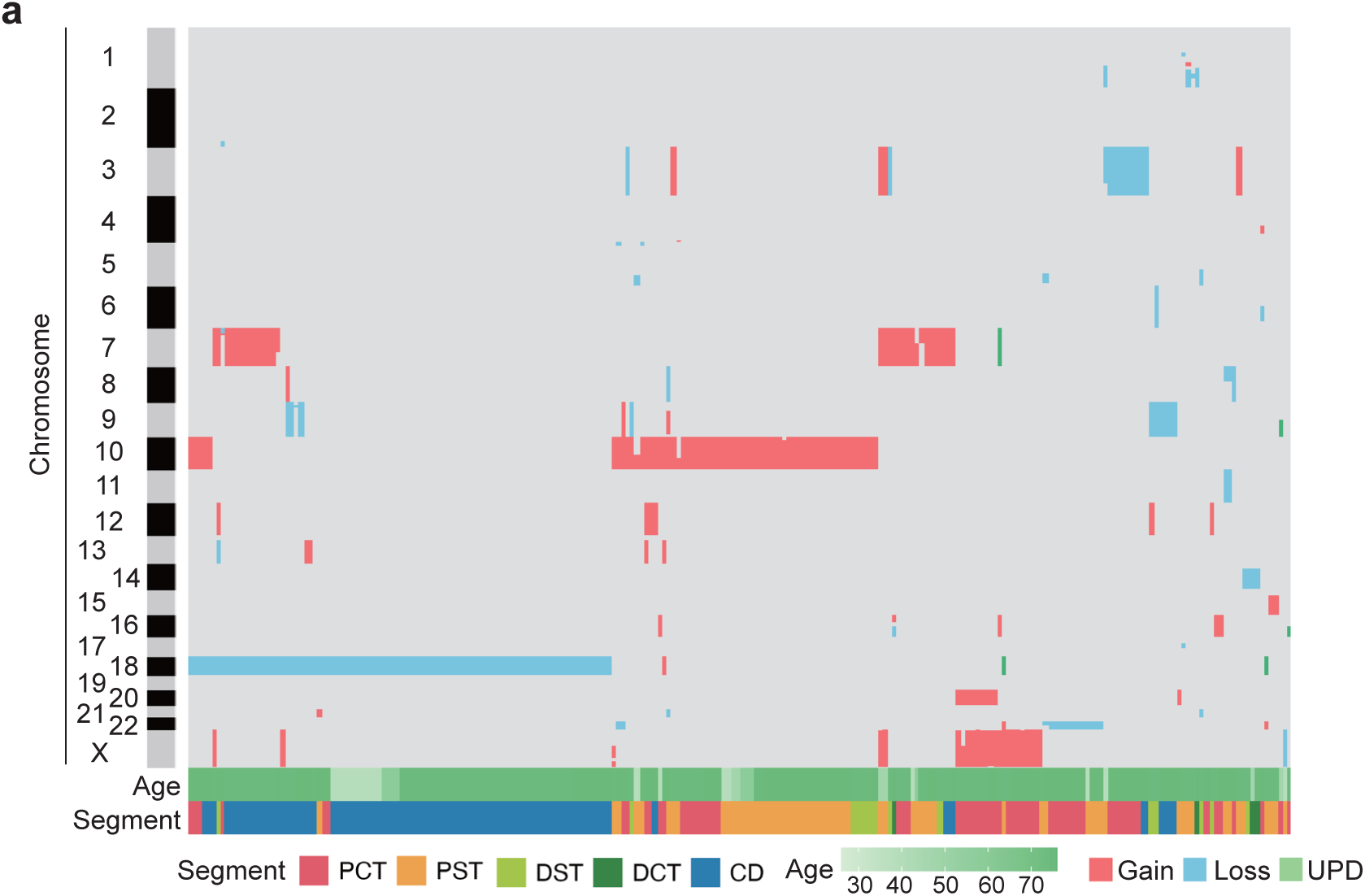
Landscape of copy number alterations (CNAs) in LMD samples. a, Landscape of CNAs in 240 of 683 LMD samples analyzed by WES. The y-axis represents chromosomes 1 through X, with heights scaled proportional to their physical lengths. Colors indicate the type of alteration (red, gain; blue, loss; green, UPD). Bottom bars represent donor age and nephron segment for each sample.

**Supplementary Figure 4.**
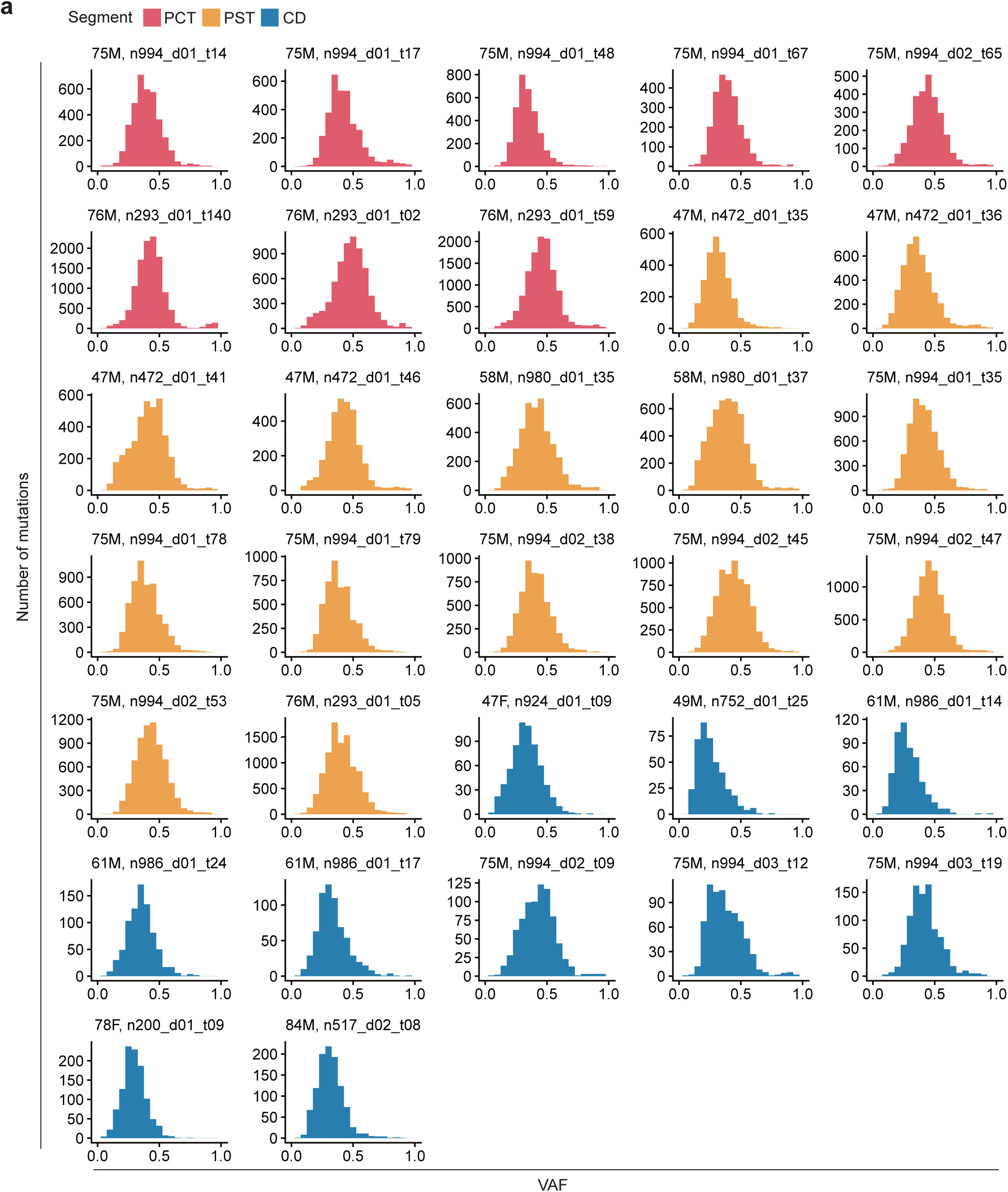
Whole genome sequencing (WGS) of LCM samples. a, Histogram of VAF of mutations detected in LCM samples using WGS. We selected a subset of LCM samples with a median VAF ≥ 0.25 in WES for WGS analysis. Samples are color-coded by segment type (PCT, red; PST, orange; CD, blue).

**Supplementary Figure 5.**
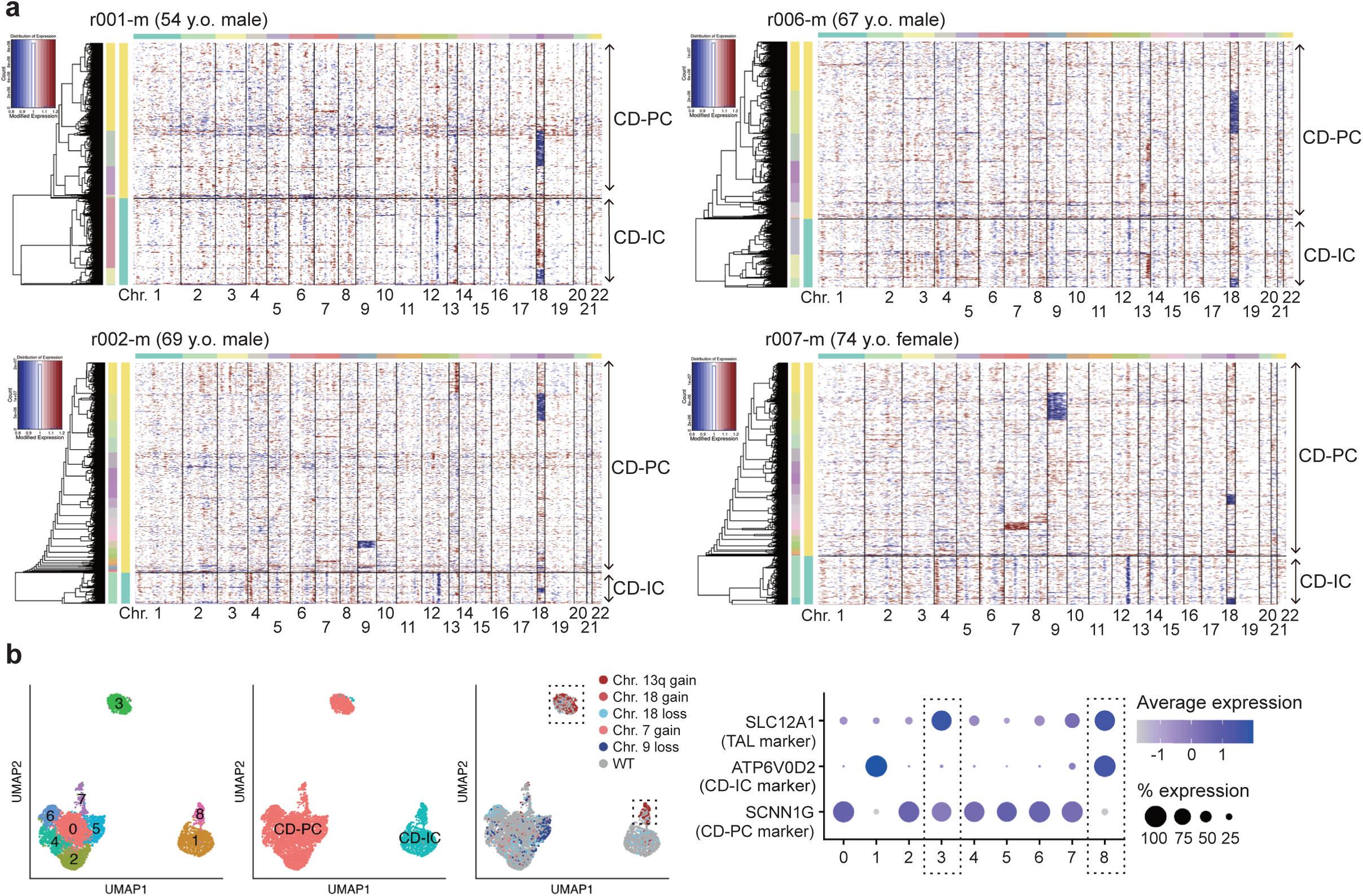
Curation process of snRNA-seq data. a, InferCNV heatmaps of pre-curated CD-PCs and CD-ICs. **b,** UMAP plot of pre-curated CD-PCs and CD-ICs from 4 donors. Dotted lines indicate suspected doublet clusters co-expressing markers of CD-PC/TAL or CD-IC/TAL.

